# Spatial modulation of RAF by RAF/MEK glue enables full-dose combination with pan-RAF inhibitor and potent RAS-mutant tumor-selective MAPK and growth inhibition

**DOI:** 10.64898/2026.07.24.740654

**Authors:** Ana Orive-Ramos, Bijaya Gaire, Christos Adamopoulos, Beau Baars, Mathieu Desaunay, Ziyue Kou, Evangelia Matenoglou, Silvia Coma, Nayeli Gutiérrez-Trejo, Kevin Mohammed, Stuart A. Aaronson, Jian Jin, Tiphaine C. Martin, Ernesto Guccione, Evripidis Gavathiotis, Jonathan A. Pachter, Poulikos I. Poulikakos

## Abstract

The clinical benefit of MAPK-targeted therapies depends on greater pathway inhibition in tumors than normal tissues. Although pan-RAF inhibitors are active in RAS-mutant cancers, combining them with MEK inhibitors requires dose reductions due to toxicity, limiting efficacy. We show the toxicity results from MEK inhibitor–mediated feedback relief, which promotes RAF activation and pan-RAF inhibitor engagement in normal cells, narrowing the therapeutic index. We further demonstrate that MEK is exclusively cytosolic, and RAF/MEK glues overcome this limitation through spatial trapping. By stabilizing cytosolic RAF–MEK complexes, RAF/MEK glues prevent feedback-driven RAF activation in normal cells while maintaining inhibition of oncogenic RAF signaling in RAS-mutant tumors, where RAF is constitutively activated at the plasma membrane. Consequently, this enables full-dose combination with pan-RAF inhibitors, resulting in deeper MAPK suppression and robust tumor regressions in RAS-mutant models. Thus, by spatially controlling wild-type effectors, drug-induced proximity can be harnessed to increase tumor selectivity of pathway-targeted therapies.

**Significance:** MAPK-targeted therapies rarely achieve durable responses in RAS-mutant cancers due to dose-limiting toxicities. We show that RAF/MEK glues, by spatially trapping RAF, can be combined with pan-RAF inhibitors at full dose, yielding tumor-selective MAPK inhibition and tumor regressions in RAS-mutant models. Thus, drug-induced proximity can be exploited for tumor-selective therapy.

## Introduction

The RAF kinases, BRAF, CRAF, and ARAF, are central effectors of the RTK/RAS/RAF/MEK/ERK (RAS/MAPK) signaling cascade, which governs cell proliferation and survival(1). Aberrant activation of this pathway, often through oncogenic mutations in KRAS, NRAS or BRAF, drives nearly one-third of human cancers(2–5). In BRAF-mutant (BRAF(MUT)) tumors, BRAF inhibitors (BRAFis) have demonstrated efficacy(6–9) and are now used clinically in combination with MEK inhibitors (MEKis)(10–12), forming a standard-of-care regimen for patients with BRAF-mutant tumors.

We previously showed that BRAFis selectively inhibit monomeric BRAF(MUT) but are ineffective in RAS-mutant (RAS(MUT)) tumors, where wild-type BRAF (BRAF(WT)) functions as a dimer(9,13–16). To target RAF dimers, next-generation inhibitors, termed dimer-selective or pan-RAF inhibitors (pan-RAFis), have been developed(17–21) and exhibit clinical activity as single agents in RAS(MUT) cancers(21). However, unlike BRAFis, pan-RAFis induce dose-limiting toxicities when combined with MEKis, requiring dose reductions that compromise efficacy(22–24). Here, we investigate the mechanistic basis for this difference in therapeutic index between BRAFi + MEKi and pan-RAFi + MEKi combinations and explore strategies to overcome this limitation in RAS(MUT) tumors.

## Results

### Pan-RAFis are more potent in RAS(MUT) compared to wild-type RAS (RAS(WT)) cells due to increased binding to the highly active wild-type RAF conformation

To identify RAFis with potential therapeutic index in RAS(MUT) context compared to RAS(WT) (normal) cells, we interrogated the DepMap drug sensitivity portal(25). Because the dataset lacks RAS(WT) normal (non-transformed) controls, we defined a surrogate RAS(WT) group comprising cancer cell lines wild-type for *RAS*, *BRAF*, *NF1*, *PIK3CA*, and *PTEN*, representing a RAS/MAPK pathway–intact baseline. The pan-RAFis belvarafenib and naporafenib (also known as Type 2 or RAF dimer-selective inhibitors) were significantly more potent in the RAS(MUT) group compared to the RAS(WT) (**Fig. 1A**). As expected, and previously shown by us(13–16,26) and others(27,28), the clinical Type 1.5 BRAF-selective inhibitor vemurafenib showed minimal growth inhibitory activity in either RAS(MUT) or RAS(WT) cells (**Supplementary Fig. S1A**). We confirmed these findings in a panel of RAS(MUT) and RAS(WT) cell lines. Treatment with a pan-RAFi (belvarafenib(21), naporafenib(29), or LY3009120(30)), more potently suppressed cell growth (**Fig. 1B; Supplementary Fig. S1B and S1C**) and MAPK signaling (**Fig. 1C; Supplementary Fig. S1D**) in RAS(MUT) compared to RAS(WT) cells. To directly test whether RAS activation underlies this differential MAPK inhibition, we used an isogenic model with doxycycline-inducible RAS(MUT) expression (**Supplementary Fig. S1E**). Induction of RAS(MUT) to near-endogenous levels enhanced MAPK signaling suppression following treatment with belvarafenib (**Fig. 1D**) or naporafenib (**Supplementary Fig. S1F**), supporting a RAS-dependent mechanism.

**Figure 1.**
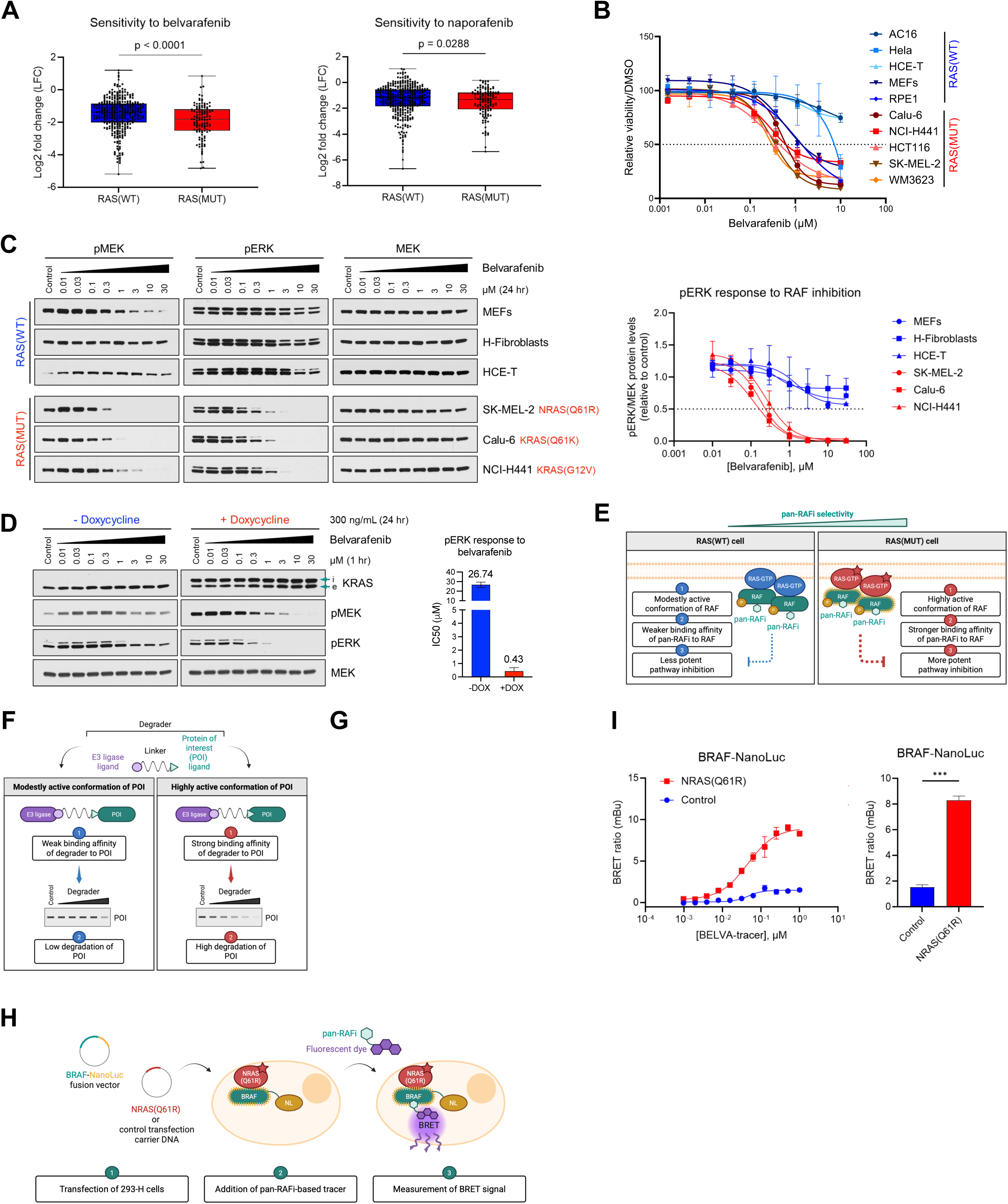
Pan-RAFis are more potent in RAS(MUT) compared to RAS(WT) cells due to increased binding to the highly active wild-type RAF conformation. **A,** Box and whisker plots show the distribution of drug response to the pan-RAFi belvarafenib and naporafenib in RAS(WT) and RAS(MUT) cells from the PRISM Repurposing Public 24Q2 dataset. *P*-values were calculated using a Mann-Whitney U test in GraphPad Prism 10. **B,** Cell growth response to belvarafenib of the indicated RAS(WT) and RAS(MUT) cell lines after 6 days of treatment. Data are represented as mean ± SEM. **C,** The indicated RAS(WT) and RAS(MUT) cell lines were treated with DMSO (control) or increasing concentrations of belvarafenib for 24 hours, and cell lysates were immunoblotted for pMEK, pERK, and MEK. The graph on the right shows the quantification of pERK normalized by the loading control MEK and relative to the control after ImageJ analysis. **D,** KRAS(Q61K) Tet-On HeLa cells were treated or not with 300 ng/mL doxycycline (DOX) for 24 hours, followed by treatment with increasing concentrations of belvarafenib for 1 hour, and cell lysates were immunoblotted for KRAS (i, induced; e, endogenous), pMEK, pERK, and MEK. The graph on the right shows the IC50 values estimated by non-linear regression from dose-response curves of pERK normalized by the loading control MEK and relative to the control after ImageJ and GraphPad Prism 10 analysis. **E,** Schematic representation showing that RAF is expressed in a highly active conformation with stronger binding to pan-RAFis in RAS(MUT) cells, whereas the conformation of RAF is modestly active with weaker binding to pan-RAFis in RAS(WT) cells. This differential binding correlates with differential MAPK signaling inhibition and therefore determines pan-RAFi selectivity. Created by Ana Orive-Ramos in BioRender. **F,** Schematic representation of a degrader and the use of degraders to evaluate degradation of a protein of interest (POI) as a surrogate of binding of a POI ligand (warhead) to the POI. Created by Ana Orive-Ramos in BioRender. **G,** RAS(WT) and RAS(MUT) cells (MEFs and HCT116, respectively) were treated with the indicated concentrations of pan-RAF-D for 18 hours, and cell lysates were immunoblotted for BRAF, CRAF, ARAF, pMEK, pERK, and MEK. **H,** Schematic representation showing the NanoBRET target engagement assay performed in 293-H cells, co-transfected with BRAF-NanoLuc and pMEV-NRAS(Q61K)-2HA or a control transfection carrier DNA and incubated with pan-RAFi-based tracers. Created by Ana Orive-Ramos in BioRender. **I,** Dose-dependent BRET signal measured in 293-H cells co-transfected as described in **H** and incubated with BELVA-tracer. The graph on the right shows BRET signal at tracer concentration of 1 µM. Data are represented as mean ± SEM, and statistical comparison was performed using a two-tailed unpaired Welch’s *t* test in GraphPad Prism 10 (***, *P* < 0.001).

Crystal structures of pan-RAFis bound to RAF show the kinase in an active αC-IN conformation(15,17,21), suggesting that these inhibitors preferentially engage active RAF. We hypothesized that in RAS(MUT) cells, RAF adopts a more active conformation that confers stronger pan-RAFi binding, whereas in RAS(WT) cells, RAF is in a less active conformation with reduced affinity (**Fig. 1E**). To test this, we applied Proteolysis Targeting Chimera (PROTAC) technology(31) to monitor endogenous target engagement via degradation (**Fig. 1F**).

We generated a pan-RAFi-based degrader (pan-RAF-D) by linking the pan-RAFi LY3009120 to a VHL1 binding moiety (**Supplementary Fig. S1G**). BRAF degradation by pan-RAF-D was prevented by treatment with proteasome or neddylation inhibitor, or excess of either VHL ligand or pan-RAFi (**Supplementary Fig. S1H**), confirming pan-RAF-D-mediated protein loss is consistent with a PROTAC mechanism of action. Treatment with pan-RAF-D induced degradation of RAF paralogs (i.e. BRAF, CRAF, and ARAF) in RAS(MUT) but not in RAS(WT) cells **(Fig. 1G, Supplementary Fig. S1I)**, associated with increased RAF dimerization in these cells (**Supplementary Fig. S1J**) and thus consistent with the extent of RAF degradation by pan-RAF-D being proportional to RAF activation state. To further establish the causal relationship between RAS activity and RAF engagement by pan-RAF-D, we employed a pharmacological approach. Treatment of RAS(MUT) cells with the RASmultiON inhibitor daraxonrasib(32,33) reduced BRAF/CRAF dimerization (**Supplementary Fig. S1K**) and attenuated pan-RAF-D-induced RAF degradation (**Supplementary Fig. S1L**). As expected, a BRAFi-based degrader (SJF-0628(34), BRAF-D) selectively degraded BRAF(V600E) in BRAF(MUT) cells, but had no effect on RAF paralogs in RAS(MUT) or RAS(WT) cells (**Supplementary Fig. S1M and S1N**), in line with previous studies by us(13–16) and others(35), establishing that BRAFis do not bind dimeric RAF.

To independently assess RAS-dependent RAF binding, we developed a NanoBRET assay using novel, cell-permeable fluorescent tracers, BELVA-tracer and LY-tracer, which we specifically designed and synthesized by covalently conjugating pan-RAFis compounds (belvarafenib or LY3009120(18)) to a fluorescent dye (**Fig. 1H; Supplementary Fig. S1O**). 293-H cells were transfected with BRAF-NanoLuc, and BRET signal was measured after incubation with tracer, with or without co-expression of RAS(MUT). Co-expression of RAS(MUT) increased binding of both BELVA-tracer (**Fig. 1I**) and LY-tracer (**Supplementary Fig. S1P**), in line with RAS activation promoting enhanced pan-RAFi binding to RAF. To further confirm this in an isogenic setting, we utilized doxycycline-inducible MEFs (**Supplementary Fig. S1Q).** In these cells, RAS(MUT) expression also increased tracer binding relative to parental cells (**Supplementary Fig. S1R**). Finally, pharmacological inhibition of RAS(MUT) with daraxonrasib reduced tracer binding in RAS(MUT)-expressing cells (**Supplementary Fig. S1S**), consistent with RAS activity directly driving enhanced pan-RAFi engagement.

### MEKis and RAF/MEK glues are more potent in RAS(MUT) over RAS(WT) cells

To assess MEKis potency in the RAS(MUT) versus RAS(WT) cellular contexts, we first analyzed the DepMap drug sensitivity portal. We found that MEKis (e.g. cobimetinib, trametinib) also show preferential growth inhibitory activity in RAS(MUT) compared to RAS(WT) cells (**Fig. 2A**). Consistently, treatment of our panel of RAS(MUT) and RAS(WT) cell lines with the MEKis trametinib(36) or cobimetinib(37) resulted in more potent suppression of cell growth (**Fig. 2B; Supplementary Fig. S2A and S2B**) and MAPK signaling (**Fig. 2C; Supplementary Fig. S2C**) in RAS(MUT) cells. Inducible RAS(MUT) expression in an isogenic model also resulted in more potent MAPK suppression by MEKi (**Fig. 2D**). Furthermore, a MEKi-based degrader (MS934(38), MEK-D) induced greater MEK degradation in RAS(MUT) over RAS(WT)-expressing cells (**Fig. 2E; Supplementary Fig. S2D**), consistent with enhanced MEKi binding in the high RAS activity state.

**Figure 2.**
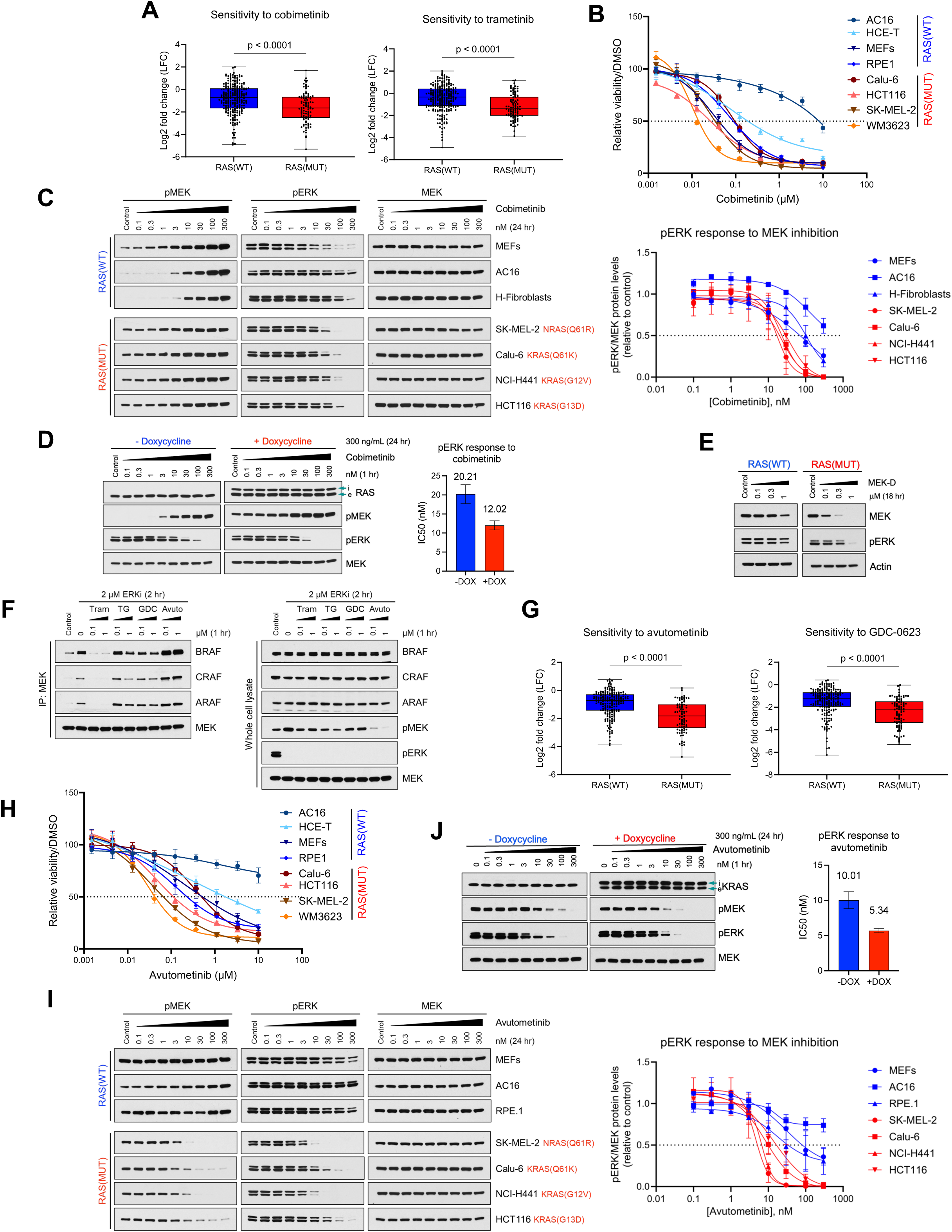
MEKis and RAF/MEK glues are more potent in RAS(MUT) over RAS(WT) cells. **A,** Box and whisker plots show the distribution of drug response to the MEKi trametinib and cobimetinib in RAS(WT) and RAS(MUT) cells from the PRISM Repurposing Public 24Q2 dataset. *P*-values were calculated using a Mann-Whitney U test in GraphPad Prism 10. **B,** Cell growth response to cobimetinib of the indicated RAS(WT) and RAS(MUT) cell lines after 6 days of treatment. Data are represented as mean ± SEM. **C,** The indicated RAS(WT) and RAS(MUT) cell lines were treated with DMSO (control) or increasing concentrations of cobimetinib for 24 hours, and cell lysates were immunoblotted for pMEK, pERK, and MEK. The graph on the right shows the quantification of pERK normalized by the loading control MEK and relative to the control after ImageJ analysis. **D,** KRAS(Q61K) Tet-On HeLa cells were treated or not with 300 ng/mL doxycycline (DOX) for 24 hours, followed by treatment with increasing concentrations of cobimetinib for 1 hour, and cell lysates were immunoblotted for RAS (i, induced; e, endogenous), pMEK, pERK, and MEK. The graph on the right shows the IC50 values estimated by non-linear regression from dose-response curves of pERK normalized by the loading control MEK and relative to the control after ImageJ and GraphPad Prism 10 analysis. **E,** RAS(WT) and RAS(MUT) cells (MEFs and HCT116, respectively) were treated with the indicated concentrations of MEK-D for 18 hours, and cell lysates were immunoblotted for MEK, pERK, and actin. **F,** RAS(MUT) cells were pre-treated with 2 μΜ of the ERK inhibitor (ERKi) SCH772984 for 2 hours, followed by treatment with the indicated concentrations of the MEK inhibitors trametinib (Tram), or RAF/MEK glues trametiglue (TG), GDC-0623 (GDC), and avutometinib (Avuto) for 1 hour. Cell lysates were either subjected to immunoprecipitation with a MEK antibody followed by immunoblotting for BRAF, CRAF, ARAF, and MEK or immunoblotted with the indicated antibodies. **G,** Box and whisker plots show the distribution of drug response to the RAF/MEK glues avutometinib and GDC-0623 in RAS(WT) and RAS(MUT) cells from the PRISM Repurposing Public 24Q2 dataset. *P*-values were calculated using a Mann-Whitney U test in GraphPad Prism 10. **H,** Cell growth response to avutometinib of the indicated RAS(WT) and RAS(MUT) cell lines after 6 days of treatment. Data are represented as mean ± SEM. **I,** The indicated RAS(WT) and RAS(MUT) cell lines were treated with DMSO (control) or increasing concentrations of avutometinib for 24 hours, and cell lysates were immunoblotted for pMEK, pERK, and MEK. The graph on the right shows the quantification of pERK normalized by the loading control MEK and relative to the control after ImageJ analysis. **J,** KRAS(Q61K) Tet-On HeLa cells were treated or not with 300 ng/mL doxycycline (DOX) for 24 hours, followed by treatment with increasing concentrations of avutometinib for 1 hour, and cell lysates were immunoblotted for KRAS (i, induced; e, endogenous), pMEK, pERK, and MEK. The graph on the right shows the IC50 values estimated by non-linear regression from dose-response curves of pERK normalized by the loading control MEK and relative to the control after ImageJ and GraphPad Prism 10 analysis.

Αvutometinib (also known as VS-6766 or CH5126766(39)) is a representative RAF/MEK glue, a class of small molecules that bind and inhibit MEK while stabilizing its interaction with RAF. Avutometinib has gained accelerated approval in combination with the FAK inhibitor defactinib for the treatment of recurrent KRAS(MUT) low-grade serous ovarian cancer(40). To isolate its direct allosteric effect on MEK from RAF-MEK complex formation due to MAPK feedback relief, we pretreated the cells with an ERK inhibitor (to relieve negative feedback), then treated with MEKi or RAF/MEK-glue and performed immunoprecipitation. As previously reported(16,41), the MEKi trametinib disrupted RAF-MEK complexes, whereas avutometinib robustly promoted their formation (**Fig. 2F**). Two other reported RAF/MEK-glues, GDC-0623(42) and trametiglue(43), did not promote or disrupt RAF-MEK binding under these conditions (**Fig. 2F**).

We next evaluated RAF/MEK glue potency in the RAS(MUT) and RAS(WT) contexts. In both the DepMap portal (**Fig. 2G)** and our panel of cell lines (**Fig. 2H; Supplementary Fig. S2B**), avutometinib and GDC-0623 were more potent in suppressing cell growth of RAS(MUT) compared to RAS(WT). Similarly, MAPK signaling suppression was more potent in RAS(MUT) cells, both in endogenous models (**Fig. 2I**) and in the RAS(MUT)-inducible setting (**Fig. 2J**).

Thus, pan-RAFis, MEKis, and RAF/MEK glues all show greater potency against RAS(MUT) over RAS(WT) cells as single agents, both in terms of growth inhibition and MAPK pathway suppression.

### Pan-RAFis selectivity for RAS(MUT) cells is abrogated by co-treatment with MEKis

Pan-RAFis have shown clinical activity in patients with RAS(MUT) tumors(21). However, unlike BRAFi + MEKi combinations, which are tolerated at the maximum tolerated dose (MTD) of each drug, pan-RAFi + MEKi combinations require significant dose reductions from each drug’s single MTD, and are associated with increased toxicities, compared to monotherapy(22–24).

To understand the molecular basis of this increased toxicity, we compared the effects of combined pan-RAFi + MEKi in RAS(MUT) and RAS(WT) cells. In RAS(MUT) cells, the combination of the pan-RAFi belvarafenib with MEKi (trametinib or cobimetinib) induced greater suppression of cell growth (**Fig. 3A**) and MAPK signaling (**Fig. 3B**) than either inhibitor alone. Unexpectedly, in RAS(WT) cells, the combination of belvarafenib (at a concentration that was inactive in RAS(WT) normal cells alone) with a low concentration of MEKi (trametinib or cobimetinib) resulted in “adverse synergy”: the combination induced dramatically greater suppression of cell growth (**Fig. 3A**) and MAPK signaling (**Fig. 3B and 3C**) than either inhibitor alone, thereby eliminating the preferential potency of these agents for the RAS(MUT) context. This adverse synergy was further confirmed in the isogenic RAS(MUT) inducible setting, where the combination of belvarafenib and MEKi similarly suppressed MAPK signaling in RAS(MUT) and RAS(WT) cells (**Supplementary Fig. S3A)**. As expected, a BRAFi (vemurafenib) did not enhance MAPK signaling inhibition in RAS(WT) cells when combined with MEKi (**Fig. 3C**), consistent with the good tolerability of BRAFi + MEKi combinations in clinical settings (10,11).

**Figure 3.**
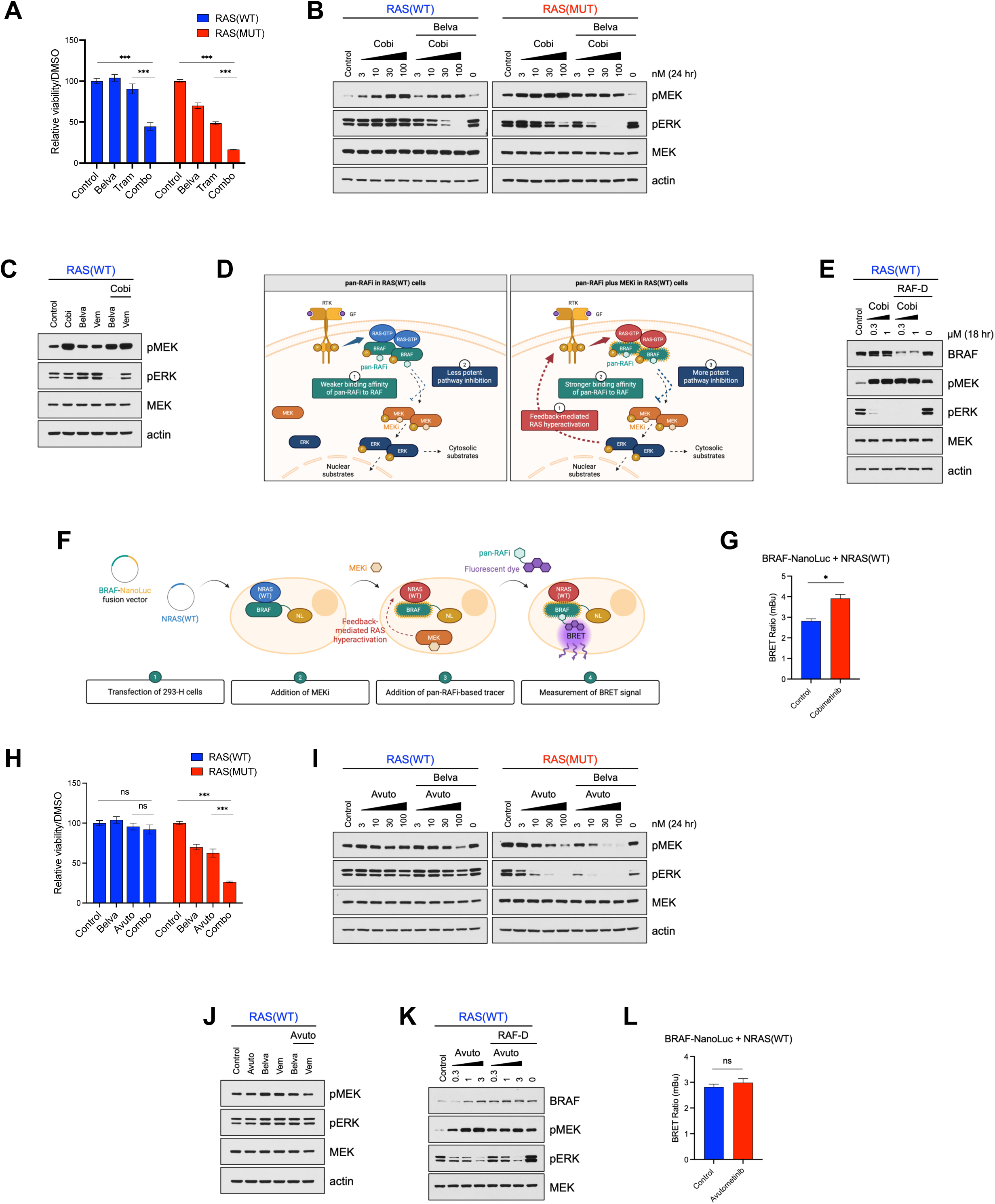
Pan-RAFis selectivity for RAS(MUT) cells is abrogated by co-treatment with MEKis, due to the relief of the negative feedback in RAS(WT) cells, but it is retained when combined with a RAF/MEK glue. **A,** Cell growth response to 250 nM belvarafenib (belva), 3 nM trametinib (tram), or the combination in RAS(WT) and RAS(MUT) cells (MEFs and SK-MEL-2 cells, respectively) after 6 days of treatment. Data are represented as mean ± SEM and statistical comparisons were performed using one way ANOVA in GraphPad Prism 10 (***, *P* < 0.001). **B,** RAS(WT) and RAS(MUT) cells (MEFs and SK-MEL-2 cells, respectively) were treated with DMSO (control), 200 nM belvarafenib (belva), or increasing concentrations of cobimetinib (cobi) alone or in combination with 200 nM belvarafenib for 24 hours, and cell lysates were immunoblotted for pMEK, pERK, MEK, and actin. **C,** MEFs were treated with DMSO (control), 30 nM cobimetinib (cobi), 200 nM belvarafenib (belva), 250 nM vemurafenib (vem), or the combinations belva + cobi or vem + cobi for 24 hours. Cell lysates were immunoblotted for pMEK, pERK, MEK, and actin. **D,** Schematic representation showing different conformations of RAF in RAS(WT) cells treated with a pan-RAFi alone or in combination with a MEKi. The addition of a MEKi promotes the highly active conformation of RAF in these cells through the relief of negative feedback, leading to better binding affinity of the pan-RAFi to RAF and subsequently more potent pathway inhibition. Created by Ana Orive-Ramos in BioRender. **E,** MEFs were treated with DMSO (control) or increasing concentrations of cobimetinib (cobi) alone or in combination with 0.5 µM pan-RAF-D for 18 hours, and cell lysates were immunoblotted for BRAF, pMEK, pERK, MEK, and actin. **F,** Schematic representation of the NanoBRET target engagement assay performed in 293-H cells, co-transfected with BRAF-NanoLuc and pMEV-NRAS(Q61K)-2HA and treated with a MEKi before incubating them with pan-RAFi-based tracers. Created by Ana Orive-Ramos in BioRender. **G,** BRET signal measured in 293-H cells co-transfected as described in **F** and treated with DMSO (control) or 3 µM cobimetinib for 2 hours before incubating them with 1 µM BELVA-tracer. Data are represented as mean ± SEM. **H,** Cell growth response to 250 nM belvarafenib (belva), 30 nM avutometinib (avuto), or the combination in RAS(WT) and RAS(MUT) cells (MEFs and SK-MEL-2 cells, respectively) after 6 days of treatment. Data are represented as mean ± SEM and statistical comparisons were performed using one way ANOVA in GraphPad Prism 10 (***, *P* < 0.001). Control and belvarafenib-treated conditions are the same as those presented in Fig. 3A. **I,** RAS(WT) and RAS(MUT) cells (MEFs and SK-MEL-2 cells, respectively) were treated with DMSO (control), 200 nM belvarafenib (belva), or increasing concentrations of avutometinib (avuto) alone or in combination with 200 nM belvarafenib for 24 hours, and cell lysates were immunoblotted for pMEK, pERK, MEK, and actin. **J,** MEFs were treated with DMSO (control), 30 nM avutometinib (avuto), 200 nM belvarafenib (belva), 250 nM vemurafenib (vem), or the combinations belva + avuto or vem + avuto for 24 hours. Cell lysates were immunoblotted for pMEK, pERK, MEK, and actin. **K,** MEFs were treated with DMSO (control) or increasing concentrations of avutometinib (avuto) alone or in combination with 0.5 µM pan-RAF-D for 18 hours, and cell lysates were immunoblotted for BRAF, pMEK, pERK, and MEK. **L,** BRET signal measured in 293-H cells co-transfected as described in **F** and treated with DMSO (control) or 3 µM avutometinib for 2 hours before incubating them with 1 µM BELVA-tracer. Data are represented as mean ± SEM. Statistical comparison in **G** and **L** was performed using a one-way ANOVA test in GraphPad Prism 10 (ns, non-significant; *, *P* < 0.05).

### Relief of negative feedback by MEKis in RAS(WT) cells promotes the activated dimeric RAF conformation, resulting in increased binding to pan-RAFis

Previous studies by us(16,44) and others(45–47) have established relief of negative feedback and upstream RTK/RAS activation as a common homeostatic response to RAS/MAPK inhibition. To explain the markedly increased potency of combined pan-RAFi + MEKi in RAS(WT) cells compared to single treatments, we hypothesized that MEKis promote the highly active RAF conformation through relief of negative feedback in RAS(WT) cells (**Fig. 3D**).

To test this, we first examined RAF conformation using the pan-RAF-D and BRAF-D degraders in RAS(WT) cells. In the absence of MEKi, BRAF expression levels were unaffected by either pan-RAF-D or BRAF-D (**Supplementary Fig. S3B**). Upon MEKi treatment, BRAF became susceptible to degradation by pan-RAF-D, but not BRAF-D (**Fig. 3E; Supplementary Fig. S3B**), consistent with MEKi-induced feedback relief promoting the highly active RAF conformation with increased binding to pan-RAF-D and its warhead. Consistent results were observed using the NanoBRET assay with the BELVA-tracer (a fluorescent derivative of belvarafenib) (**Fig. 3F**). Treatment with the MEKi cobimetinib increased BRET signal (**Fig. 3G**), supporting the model that MEKi-mediated feedback relief enhances RAS activation, thereby promoting an active RAF conformation with stronger tracer binding.

Together, these data show that pan-RAFi + MEKi combination therapy exhibits potentiated toxicity in RAS(WT) cells, markedly exceeding the additive effects of the individual agents. This potentiated toxicity arises because MEKi relieves feedback inhibition, promoting the highly active RAF conformation that binds pan-RAFi more effectively.

### A pan-RAFi + RAF/MEK glue combination retains tumor-selective potency in RAS(MUT) over RAS(WT) cells

To identify pan-RAFi-based combinations that preserve selectivity for RAS(MUT) cells, thereby enabling full dosing of both drugs to improve efficacy, we evaluated the combination of the pan-RAFi belvarafenib with the RAF/MEK glue avutometinib. This combination suppressed both cell growth (**Fig. 3H**) and MAPK signaling (**Fig. 3I**) more potently than either agent alone in RAS(MUT) cells. Strikingly, unlike pan-RAFi + MEKi, the belvarafenib + avutometinib combination retained tumor cell-selective potency, as evidenced by more potent cell growth and MAPK signaling suppression in RAS(MUT) over RAS(WT) cells (**Fig. 3H and 3I**) and minimal MAPK signaling inhibition in RAS(WT) cells (**Fig. 3J**). We further confirmed this RAS(MUT)-selective suppression of MAPK signaling in an isogenic RAS(MUT) inducible setting, where belvarafenib combined with avutometinib potently inhibited MAPK signaling in RAS(MUT) cells while sparing RAS(WT) cells (**Supplementary Fig. S3C)**. A similar pattern of RAS(MUT)-selective MAPK suppression was observed with another pan-RAFi + RAF/MEK glue combination (naporafenib + avutometinib), whereas naporafenib combined with the MEKi trametinib showed comparable MAPK inhibition in RAS(MUT) and RAS(WT) cells (**Supplementary Fig. S3D**).

To further validate the model using orthogonal approaches with independent readouts, we assessed whether avutometinib alters RAF conformation or binding in RAS(WT) cells by monitoring BRAF degradation using pan-RAF-D and BRET signal induced by BELVA-tracer in the NanoBRET assay. First, unlike MEKi, avutometinib did not promote BRAF degradation by pan-RAF-D (**Fig. 3K**), indicating that it does not induce a high-affinity RAF conformation in normal cells. Second, avutometinib did not increase BRET signal induced by BELVA-tracer binding to BRAF-NanoLuc (**Fig. 3L**), further supporting that, in contrast to MEKis, RAF/MEK glues do not promote the active RAF conformation in RAS(WT) cells. We next evaluated the belvarafenib and avutometinib combination in a panel of KRAS G12D mutant cell lines and found that growth suppression by the combination was associated with KRAS-dependency (**Supplementary Fig. S3E and S3F**). In cell lines highly dependent on KRAS (AsPC-1 and HPAFII), the combination of belvarafenib and avutometinib showed synergy effects. In contrast, in cell lines less dependent on KRAS (SW1990 and PANC-1), the additive effect of the drugs was attenuated. Finally, we assessed the efficacy of this combination in models of acquired resistance to RAS inhibitors: SK-MEL-2 cells that have developed resistance to daraxonrasib and ASPC1 cells that have developed resistance to the KRASG12D inhibitor MRTX-1133 (48). Belvarafenib plus avutometinib was similarly potent in suppressing cell growth and MAPK signaling in both parental and KRASG12D inhibitor-resistant cells (**Supplementary Fig. S3G-J**).

Together, these data indicate that the pan-RAFi plus RAF/MEK glue combination can be an effective therapeutic strategy for both treatment-naïve RAS(MUT) tumors and those that have developed resistance to RAS inhibitors, a scenario of increased clinical relevance.

### MEK is constitutively cytosolic, and the RAF/MEK glue promotes RAF inactivation by trapping RAF with MEK in the cytosol

Unlike the pan-RAFi + MEKi combination, pan-RAFi + RAF/MEK glue retains selective potency for RAS(MUT) over RAS(WT) cells. To explain this difference, we first compared how MEKi and RAF/MEK glue affect RAS activation and RAF complex formation in RAS(WT) cells. Treatment with either MEKi (cobimetinib) or RAF/MEK glue (avutometinib) promoted similar levels of feedback-induced RAS activation (**Fig. 4A**), indicating that their differential effects occur downstream of RAS. As expected, in RAS(WT) cells, MEKi induced BRAF-CRAF complex formation (**Fig. 4B; Supplementary Fig. S4A**), consistent with enhanced RAF activation following feedback relief. In contrast, RAF/MEK glues promoted MEK-BRAF complex formation while preventing BRAF-CRAF dimerization (**Fig. 4B; Supplementary Fig. S4A**).

**Figure 4.**
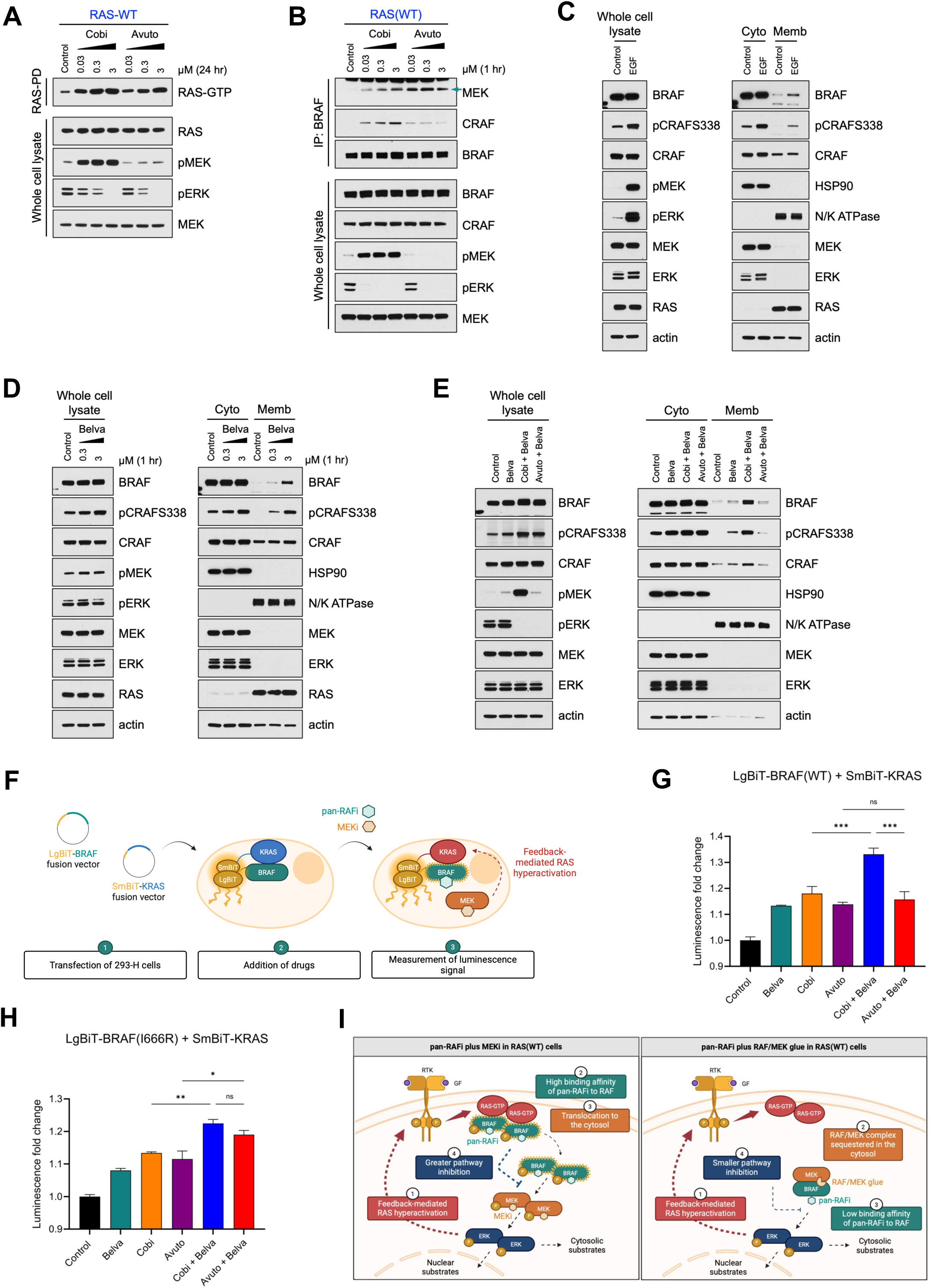
MEK is constitutively cytosolic, and the RAF/MEK glue promotes RAF inactivation by trapping RAF in the cytosol. **A,** MEFs were treated with increasing concentrations of cobimetinib (cobi) or avutometinib (avuto) for 24 hours. Cell lysates underwent a RAS-GTP pull-down assay, followed by immunoblotting with a RAS antibody, or were immunoblotted with the indicated antibodies. **B,** MEFs were treated with increasing concentrations of cobimetinib (cobi) or avutometinib (avuto) for 1 hour. Cell lysates were subjected to immunoprecipitation with a BRAF antibody, followed by immunoblotting for MEK, CRAF, and BRAF, or were immunoblotted with the indicated antibodies. **C,** HeLa cells were serum-starved overnight and treated with DMSO (control) or 50 ng EGF for 5 minutes. Cells underwent cell fractionation, and cytosolic (cyto) and membrane (memb) fractions, alongside whole cell lysates, were immunoblotted with the indicated antibodies. **D,** HeLa cells were treated with increasing concentrations of belvarafenib (belva) for 1 hour. Cells underwent cell fractionation, and cytosolic (cyto) and membrane (memb) fractions, along with whole cell lysates, were immunoblotted with the indicated antibodies. **E,** HeLa cells were treated with 0.3 µM belvarafenib (belva) for 1 hour or with 3 µM cobimetinib (cobi) or 3 µM avutometinib (avuto) for 1 hour, followed by treatment with 0.3 µM belvarafenib for 1 hour. Cells underwent cell fractionation, and cytosolic (cyto) and membrane (memb) fractions, along with whole cell lysates, were immunoblotted with the indicated antibodies. **F,** Schematic representation of the RAF-RAS NanoBiT protein-protein interaction assay. 293-H cells were co-transfected with LgBiT-BRAF and SmBiT-KRAS, and were treated with a MEKi or RAF/MEK glue (not shown), followed by a pan-RAFi, before measuring luminescence. Created by Ana Orive-Ramos in BioRender. **G,** Luminescence signal fold change measured in 293-H cells co-transfected as in **F** and treated with 1 µM belvarafenib (belva) for 1 hour, 3 µM cobimetinib (cobi) for 2 hours, 3 µM avutometinib (avuto) for 2 hours, or with 3 µM cobimetinib or 3 µM avutometinib for 1 hour, followed by treatment with 1 µM belvarafenib for 1 hour. Data are represented as mean ± SEM. Statistical comparison was performed using a one-way ANOVA test in GraphPad Prism 10 (ns, non-significant; ***, *P* < 0.001). **H,** Luminescence signal fold change measured in 293-H cells co-transfected as in **F** with LgBiT-BRAF(I666R) and SmBiT-KRAS and treated with 1 µM belvarafenib (belva) for 1 hour, 3 µM cobimetinib (cobi) for 2 hours, 3 µM avutometinib (avuto) for 2 hours, or with 3 µM cobimetinib or 3 µM avutometinib for 1 hour, followed by treatment with 1 µM belvarafenib for 1 hour. Data are represented as mean ± SEM. Statistical comparison was performed using a one-way ANOVA test in GraphPad Prism 10 (ns, non-significant; *, *P* < 0.05; **, *P* < 0.01). **I,** Schematic representation showing that MEK is constitutively cytosolic, and while both MEKi and RAF/MEK glue promote feedback-mediated RAS hyperactivation, the RAF/MEK glue sequesters RAF in the cytosol, resulting in RAF inactivation, which leads to low binding of pan-RAFi to RAF and reduced pathway inhibition when combined with pan-RAFis. Created by Ana Orive-Ramos in BioRender.

RAF inactivation by RAF/MEK glues has previously been proposed based on structural studies (49,50). Still, the underlying mechanism remains unresolved, as other structural studies have captured RAF/MEK complexes in both inactive and active conformations(51). These findings suggest that RAF inactivation by RAF/MEK glue does not arise from a conformational (allosteric) mechanism, but may instead involve changes in subcellular localization or altered protein interactions.

RAF activation in cells requires translocation from the cytosol to the cell membrane(52), RAS binding, and dimerization(53). To identify the step(s) disrupted by RAF/MEK glue, we performed subcellular fractionation. As expected, in RAS(WT) cells, EGF stimulation or treatment with a pan-RAFi (belvarafenib or naporafenib) promoted BRAF and CRAF recruitment to the plasma membrane (**Fig. 4C and 4D; Supplementary Fig. S4B**). Surprisingly, MEK remained cytosolic under all conditions (**Fig. 4C and 4D; Supplementary Fig. S4B**), suggesting that tethering RAF to MEK may block RAF translocation and activation. Consistent with this, in RAS(WT) cells, the pan-RAFi + MEKi combination increased RAF membrane recruitment, while the pan-RAFi + RAF/MEK glue combination completely blocked it (**Fig. 4E**). Similar results were obtained using trametinib versus NST-628(51), another RAF/MEK glue in clinical development, in combination with pan-RAFi (**Supplementary Fig. S4C**). These findings indicate that in RAS(WT) cells, RAF cytosolic trapping is a general feature of RAF/MEK glues and is not shared by conventional MEKis.

To further test this model using an orthogonal approach, we performed an in-cell RAF-RAS NanoBit protein-protein interaction assay (**Fig. 4F**). Consistent with the cell fractionation results, the combination of a pan-RAFi with a MEKi markedly increased luminescence, indicating enhanced RAF-RAS interaction. In contrast, the pan-RAFi + RAF/MEK glue combination suppressed this increase (**Fig. 4G; Supplementary Fig. S4D**). To confirm that the cytosolic trapping is dependent on direct RAF-MEK engagement, we generated a construct expressing BRAF(I666R), a mutation that has been shown to disable BRAF-MEK binding(54). In cells expressing BRAF(I666R), the pan-RAFi + RAF/MEK glue combination lost its ability to suppress the pan-RAFi-induced enhancement of RAF-RAS interaction (**Fig. 4H**), confirming that direct BRAF-MEK binding is required for the cytosolic trapping mechanism.

Together, these results support a biochemical mechanism of action of RAF/MEK glues on RAF, in which MEK is exclusively cytosolic, and RAF/MEK glues inactivate RAF by tethering it to MEK in the cytosol, thereby preventing RAF membrane localization and its interaction with RAS (**Fig. 4I**).

### RAF/MEK glue converts cytostatic responses into robust regressions in RAS(MUT) tumors

To assess the efficacy and tolerability of combining pan-RAFi with a RAF/MEK glue *in vivo*, we first evaluated the NRAS(MUT) melanoma xenograft SK-MEL-2, a model known to be relatively sensitive to MAPK inhibition. Belvarafenib + avutometinib induced deep regressions (>30% tumor regression from baseline) in 100% of tumors (100% overall response rate [ORR]) (**Fig. 5A and 5B; Supplementary Fig. S5A**) at doses that were well tolerated without weight loss (**Supplementary Fig. S5B**). Similarly, in the KRAS(MUT) colorectal cancer xenograft LS513, which also exhibits measurable drug sensitivity, the combination produced tumor regressions in the majority of mice, with an ORR of 75% (**Fig. 5C and 5D; Supplementary Fig. S5C**), again without evidence of toxicity (**Supplementary Fig. S5D**).

**Figure 5.**
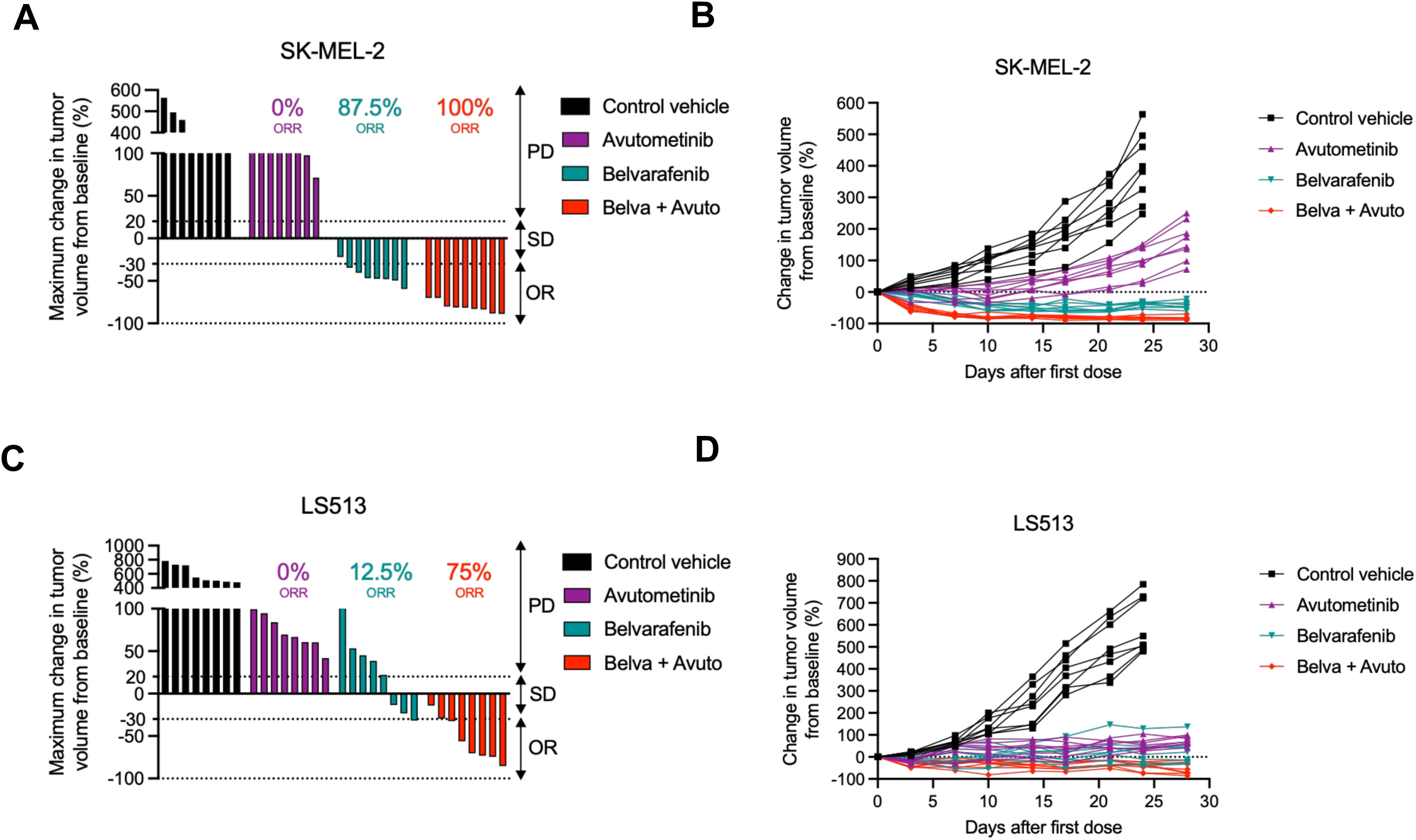
The pan-RAFi + RAF/MEK glue combination results in deep tumor regressions in RAS(MUT) tumors *in vivo*. **A,** Waterfall plot showing tumor objective response rate (ORR) in female BALB/c nude mice bearing SK-MEL-2 xenografts and treated with (i) control vehicle, (ii) avutometinib 0.1 mg/kg for 5 days on (QDx5), 2 days off, administered orally (PO), (iii) belvarafenib 15 mg/kg once daily (QD) PO, or (iv) belvarafenib 15 mg/kg QD PO combined with avutometinib 0.1 mg/kg QDx5, 2 days off, PO, for 28 days. PD, progressive disease; SD, stable disease; OR, objective response. **B,** Spider plot of change in tumor volume from baseline over time from **A**. **C,** Waterfall plot illustrating ORR in female BALB/c nude mice bearing LS513 xenografts and treated as in **A** for 28 days. PD, progressive disease; SD, stable disease; OR, objective response. **D,** Spider plot of change in tumor volume from baseline over time from **C**.

However, such sensitive models can overestimate the potential of RAS/MAPK inhibitors, since the majority of RAS(MUT) tumors in both preclinical studies and clinical trials achieve at best disease stabilization. This “therapeutic ceiling” has defined the therapeutic limitation of all current RAS/MAPK-targeted strategies, including pan-RAFi + MEKi combinations, and has prevented their clinical translation. To rigorously test whether RAF/MEK glues can overcome this barrier, we turned to NCI-H2122 cells, a KRAS(MUT) lung cancer xenograft previously reported to yield only stasis with pan-RAFi + MEKi(21). Consistent with prior reports(21), belvarafenib + cobimetinib produced only stable disease: 6/8 mice showed stable disease, and 2/8 exhibited tumor progression, resulting in a 0% ORR (**Fig. 6A and 6B**). Strikingly, replacing the MEKi with full-dose avutometinib transformed this stable disease profile into robust regressions, with 9/10 mice exhibiting ≥30% tumor shrinkage from baseline (90% ORR) (**Fig. 6A and 6B**). Pharmacodynamic analysis confirmed that tumor regressions were associated with more effective MAPK suppression: phospho-ERK immunohistochemistry demonstrated marked pathway inhibition (**Fig. 6C; Supplementary Fig. S6A**), RNA-seq analysis showed reduced MAPK transcriptional output(55,56) (**Fig. 6D**), and Western blot analysis showed markedly lower levels of pERK, as well as the downstream effectors DUSP4, DUSP6, and cyclin D1(55) (**Fig. 6E**) relative to the pan-RAFi + MEKi arm. Both regimens were well tolerated, with no weight loss or other signs of toxicity (**Supplementary Fig. S6B**).

**Figure 6.**
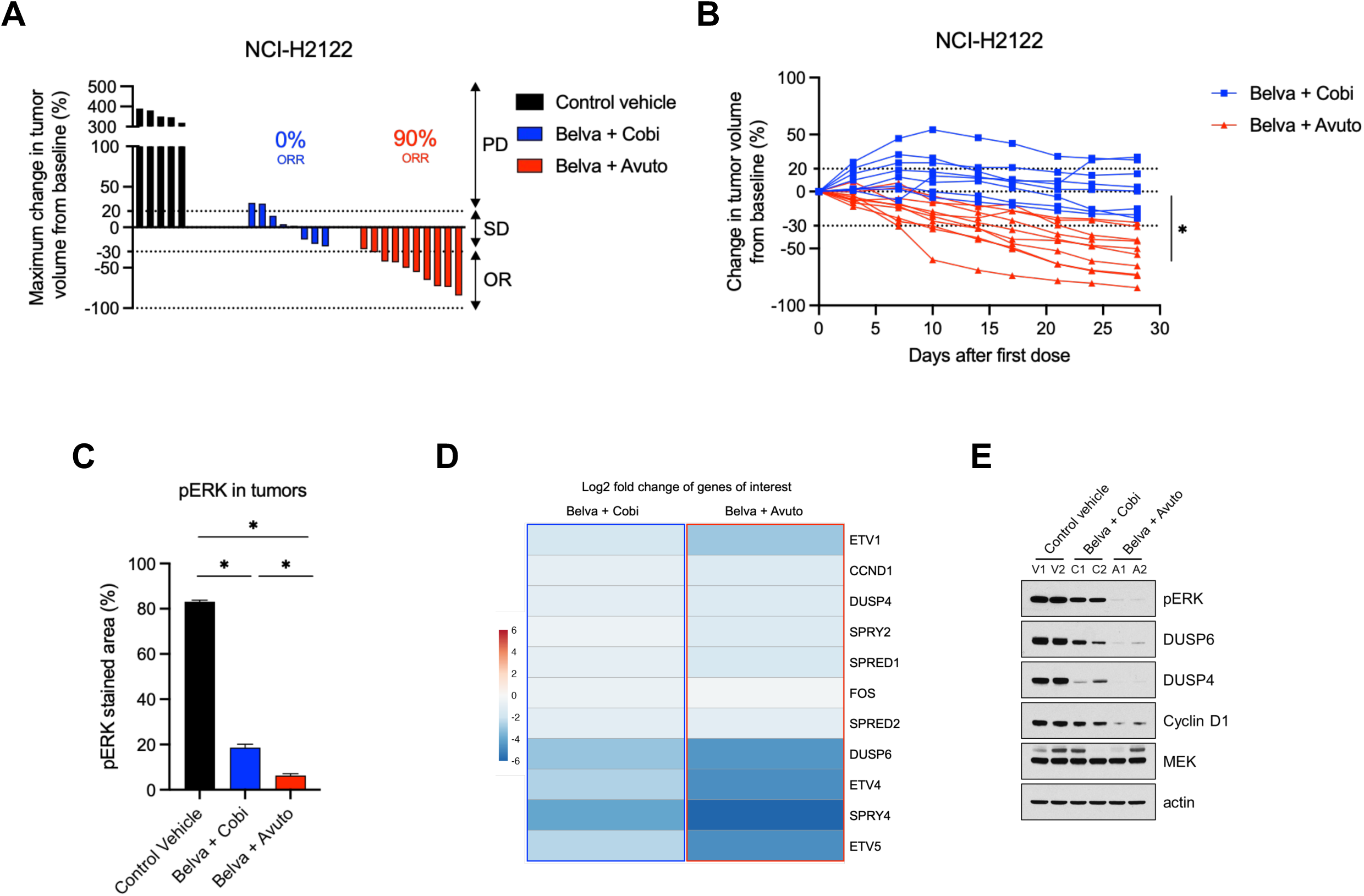
The pan-RAFi + RAF/MEK glue combination enables full dosing of each inhibitor and promotes deeper RAS(MUT) tumor regressions *in vivo* compared to pan-RAFi + MEKi. **A,** Waterfall plot showing tumor objective response rate (ORR) in female athymic Nude-*Foxn1^nu^* mice bearing NCI-H2122 xenografts and treated with (i) control vehicle, (ii) belvarafenib 15 mg/kg once daily (QD), administered orally (PO) combined with cobimetinib 5 mg/kg QD PO, or (iii) belvarafenib 15 mg/kg QD PO combined with avutometinib 0.3 mg/kg QD PO, for 28 days. PD, progressive disease; SD, stable disease; OR, objective response. **B,** Spider plot of change in tumor volume from baseline over time from **A**. Statistical comparisons were performed using a two-tailed unpaired Welch’s *t* test in GraphPad Prism 10 (*, *P* < 0.05). **C,** Quantification of the immunohistochemistry (IHC)-stained tumor sections against phosphorylated ERK (pERK) from female athymic Nude-*Foxn1^nu^* mice bearing NCI-H2122 xenografts and treated as in **A** for 3 days. Data are represented as mean ± SEM. Statistical comparison was performed using a one-way ANOVA test in GraphPad Prism 10 (*, *P* < 0.05). **D,** Heatmap of the log2 fold change expression levels of the genes of interest (MAPK effector targets) after bulk RNA sequencing analysis in tumors from female athymic Nude-*Foxn1^nu^* mice bearing NCI-H2122 xenografts and treated as in **A** for 3 days. **E,** Western blot showing expression levels of pERK, DUSP6, DUSP4, cyclin D1, MEK, and actin in tumors from female athymic Nude-*Foxn1^nu^* mice bearing NCI-H2122 xenografts and treated as in **A** for 3 days.

Together, these results demonstrate that RAF/MEK glues enable full-dose combination therapy with pan-RAFi, overcoming the “therapeutic ceiling” of MAPK-targeting regimens and producing tumor regressions even in therapy-refractory RAS(MUT) tumor models.

## Discussion

The clinical success of BRAF plus MEK inhibitor combinations in BRAF(MUT) melanomas and other BRAF(MUT) cancers inspired efforts to apply a similar approach to other RAS/MAPK-driven cancers, particularly RAS(MUT) cancers, which collectively account for about a third of human malignancies. Despite the initial promise of pan-RAFis as single agents, which were developed to inhibit RAF dimers, their combinations with MEKis have led to dose-limiting toxicities and reduced efficacy. Here, we show that the activity of pan-RAFis as single agents stems from their preferential binding to highly activated RAF in RAS(MUT) tumor cells, compared to the modestly activated RAF present in normal cells. However, when combined with MEKis, ERK-dependent negative feedback is relieved, triggering RAS activation and promoting the active RAF conformation in normal tissues. This, in turn, enhances pan-RAFi binding and leads to potent MAPK suppression in normal cells, thereby compromising the therapeutic index.

Several small molecules that both inhibit MEK and stabilize its interaction with RAF, termed RAF/MEK glues, are now approved(40) and in clinical development(39,51) (NCT05585320, NCT06326411). Although these agents have been shown to inactivate RAF in cells, the mechanism has remained unclear. Structural studies of RAF/MEK-glue-bound complexes have captured RAF in both active and inactive conformations(51), suggesting that their suppressive effect does not derive from a conformational (allosteric) mechanism. Instead, our findings reveal a spatial mechanism: RAF/MEK glues trap RAF in the cytosol by stabilizing its complex with MEK, which is constitutively cytosolic. This prevents RAF from undergoing translocation to the plasma membrane, a step required for its activation. This model explains the biochemical mechanism of RAF/MEK glue action and reframes RAS/MAPK signaling as a dynamic and spatially ordered cycle, requiring RAF to shuttle between membrane-localized activation and cytosolic engagement with downstream targets such as MEK. This aligns with prior observations of cytosolic RAF/MEK complexes as part of the normal activation-inactivation cycle of MAPK signaling(42,54).

This spatial trapping mechanism also explains the ability of RAF/MEK glues to prevent aberrant RAF activation in normal RAS (WT) cells, thereby preserving the tumor selectivity of pan-RAFi combinations. When RAF activation is driven by oncogenic RAS, as in RAS(MUT) tumors, RAF is already membrane-localized and active, allowing pan-RAFis to bind and inhibit it effectively. In normal cells, however, co-treatment with RAF/MEK glue sequesters RAF in the cytosol, reducing its activation and thereby preventing effective pan-RAFi engagement. This buffering of normal tissue sensitivity represents a distinct advantage over MEKi.

RAS(MUT) cancers remain a critical unmet clinical need. Current RAS/MAPK-directed therapies, including pan-RAFi + MEKi, are constrained by a “therapeutic ceiling”: at tolerated doses, they achieve, at best, disease stabilization rather than durable tumor regression, for the majority of patients. This limitation has prevented the clinical translation of otherwise mechanistically rational strategies. Our findings indicate that the strategy of substituting a RAF/MEK glue for a MEKi will not only alleviate dose-limiting toxicity but also promote deep regressions in refractory RAS(MUT) models.

The clinical utility of this combination spans several distinct settings. Current RAS inhibitors are restricted to specific KRAS alleles (KRAS-G12C, KRAS-G12D, or pan-KRAS G12 inhibitors such as daraxonrasib), leaving a substantial population of RAS-mutant cancers without direct RAS targeting options. Tumors harboring KRAS-G13, NRAS-Q61, or other non-G12 alleles, including NRAS-mutant melanoma for which we present data here, represent an immediate candidate population. Moreover, because our mechanism of tumor selectivity is fundamentally dependent on elevated RAS activity to engage RAF in its active conformation, highly active RAS alleles such as Q61 or G13D mutants, which drive exceptionally high RAS-GTP levels relative to G12 mutants, are particularly well-suited candidates. Beyond first-line application, acquired resistance to RAS inhibitors is frequently associated with RAS-mutant protein overexpression(57–60) and consequently RAS signaling hyperactivation, a feature that our combination exploits to establish tumor selectivity. This creates a compelling therapeutic logic: RAS inhibitor-resistance mechanisms enhance the tumor-selective therapeutic window of the pan-RAFi + RAF/MEK glue combination, positioning it as a rational post-RAS inhibitor strategy. Our data in acquired RAS inhibitor-resistant models are consistent with this, though further *in vivo* studies will be needed to fully establish the combination’s utility in the post-RAS inhibitor setting.

By leveraging complementary biochemical mechanisms: pan-RAFis to suppress RAF dimers and RAF/MEK glues to trap RAF in inactive cytosolic complexes in normal cells, this combination delivers full-dose, well-tolerated MAPK pathway suppression in a RAS(MUT) selective manner. These data provide a compelling rationale for the clinical evaluation of pan-RAFi + RAF/MEK glue combinations in RAS(MUT cancers and establish a broader conceptual advance: that drug-induced proximity(61) can be harnessed to reprogram the spatial and biochemical state of wild-type effectors in a tumor-selective fashion, thereby overcoming the therapeutic limitations of conventional targeted strategies.

## Acknowledgments

P.I.P. was supported by the US National Institutes of Health (NIH) (R01 CA285713, R01 CA240362, R01CA238229), a V Foundation Translational Grant, the Melanoma Research Alliance, the Breast Cancer Alliance, the Melanoma Research Foundation, the Irma T. Hirschl Trust and Tisch Cancer Institute developmental awards. E.G. was supported by NIH grants R01CA223243 and P30CA013330.

## MATERIALS AND METHODS

### Large-scale genomics and drug sensitivity analysis

The PRISM Repurposing Public 24Q2 dataset (https://doi.org/10.6084/m9.figshare.25917643.v1) from the Dependency Map (DepMap) portal was used to collect the log2 fold change viability values of all assessed drugs targeting components of the RAS/MAPK signaling pathway and to extract the somatic mutations of each cell line. Cell lines were annotated into two groups based on their mutational status. We considered the cell lines to be wild-type RAS (RAS(WT)) if they do not harbor any single or multiple pathogenic mutations in *KRAS*, *NRAS*, *HRAS*, *BRAF*, *PIK3CA*, *PTEN*, or *NF1*, and RAS-mutated (RAS(MUT)) if they harbor a single RAS mutation in *KRAS*, NRAS, or HRAS. To test whether the status of RAS mutation correlates with drug sensitivity, a Mann-Whitney U test (equivalent to the two-sided Wilcoxon rank sum test) between the log2 fold change viability values of RAS(WT) and RAS(MUT) cell lines was performed for each drug and visualized using GraphPad Prism 10.

### Compounds

Belvarafenib (S8853), naporafenib (S8745), LY3009120 (S7842), cobimetinib (S8041), trametinib (S2673), GDC-0623 (S7553), NST-628 (E1965), MDK7526/VHL ligand 1 (S0097), and bortezomib (S1013) were obtained from Selleckchem. MLN4924 was purchased from Cayman Chemical (15217). Daraxonrasib (HY-148439) and MRTX-1133 (HY-134813) were obtained from MedchemExpress. SJF-0628/BRAF-D (7463) was purchased from Tocris Bioscience. MS934/MEK-D was reported previously (*24*). Trametiglue was a gift from Arvin Dar (Memorial Sloan Kettering Cancer Center). Avutometinib was provided by Verastem Oncology.Compounds were dissolved in DMSO (Thermo Fisher Scientific, BP231-100) to yield a stock solution of 10 mM. Animal-free recombinant human EGF was purchased from PeproTech (AF-100-15) and reconstituted using distilled water to a concentration of 100 μg/mL.

### Cell lines

Primary dermal fibroblasts (H-Fibroblasts, PCS-201-012), SK-MEL-2 (HTB-68), Calu-6 (HTB-56), HCT116 (CCL-247), NCI-H441 (HTB-174), A-375 (CRL-1619), ASPC1 (CRL-1682), HPAFII (CRL-1997), SW1990 (CRL-2172), PANC-1 (CRL-1469), NCI-H2122 (CRL-5985), and LS513 (CRL-2134) cell lines were purchased from the American Type Culture Collection (ATCC). 293-H (11631017) and 293-FT (R70007) cells were purchased from Thermo Fisher Scientific. Mouse embryonic fibroblasts (MEFs) have been described previously (ref 8). RPE.1, AC16, and HCT-E cell lines were a generous gift from Anthony Faber (Virginia Commonwealth University). WM3623 cells and HeLa cells were kindly provided by Meenhard Herlyn (The Wistar Institute) and Ramon Parsons (Icahn School of Medicine at Mount Sinai, respectively. KRAS(Q61K) Tet-On HeLa cells and KRAS(Q61K) Tet-On MEFs were generated as described below. All the cells were cultured in DMEM (Thermo Fisher Scientific, 10313039), RPMI 1640 (Thermo Fisher Scientific, 21870092), or DMEM/F12 (Thermo Fisher Sceintific, 11320033) supplemented with 10% FBS (Biowest, S1480), 2 mM glutamine (Thermo Fisher Scientific, 25030164), 100 U/mL penicillin (Thermo Fisher Scientific, 15140163), and 100 μg/mL streptomycin (Thermo Fisher Scientific, 15140163). Cells were maintained in a humidified incubator at 37°C with 5% CO2 and were passaged 5 to 8 times. Cell lines were authenticated by LabCorp using short tandem repeat DNA profiling and were regularly tested negatively for mycoplasma using the Universal Mycoplasma Detection Kit (ATCC, 30-1012K).

### Doxycycline-inducible expression in HeLa cells and MEFs

The Tet-On system was used to conditionally express KRAS(Q61K) in HeLa cells and MEFs that were transduced with lentiviral particles. For lentiviral production, 293-FT cells were co-transfected with 1.5 μg of the packaging plasmid psPAX2 (Addgene, #12260), 4.8 μg of the envelope plasmid pMD2.G (Addgene, #12259), and 6 μg of the pTRI-BlaS transfer plasmid containing KRAS(Q61K)-2HA using Lipofectamine 2000 (Invitrogen, 11668019). pTRI-BlaS was a gift from Stuart Aaronson (Icahn School of Medicine at Mount Sinai), and KRAS(Q61K)-2HA was cloned into the plasmid by restriction enzyme digestion. After 24 hours, the 293-FT medium was replaced with fresh medium, and 48 hours after co-transfection, the medium containing lentiviral particles was collected, filtered through a 0.45 μm filter unit (Millipore Sigma SLHPR33RS), and used to infect HeLa cells or MEFs after the addition of 10 μg/mL polybrene (Sigma-Aldrich, TR-1003). HeLa cells and MEFs were cultured for 48 hours to allow for efficient selectable marker gene expression and were then selected with 3 μg/mL blasticidin (Invivogen, ant-bl-05). After a week of selection, titration of doxycycline (Millipore Sigma, D3072) allowed the expression of mutant KRAS and was evaluated by Western blot as described below.

### Generation of isogenic acquired resistant models

SK-MEL-2 daraxonrasib-acquired resistant cells (SK-MEL-2 RES) and ASPC1 MRTX1133-acquired resistant cells (ASPC1 RES) were generated by continuous dose escalation. Cells were initially exposed to low drug concentrations (daraxonrasib at 1 nM and MRTX1133 at 10 nM), which were gradually increased to high concentration (daraxonrasib at 10 nM and MRTX1133 at 100 nM). After resistance was confirmed, resistant cell lines were maintained in culture medium with corresponding drugs. Prior to experimental assays, resistant cells were cultured in drug-free medium for 3-5 days.

### Cell viability

Cells were plated in 96-well plates (Corning, 3595) at a density of 0.25-1 × 10^3^ cells per well. The next day, cells were treated with inhibitors as indicated in regular growth media for 6 days. Growth media with or without inhibitors was replaced after 3 days of treatment. Cells were incubated with the water-soluble tetrazolium salt Cell Counting kit-8 solution (Dojindo Laboratories, CK04-11) for 90 minutes at 37°C according to the manufacturer’s recommendations, and the absorbance was read at 450 nM using the GloMax® Discover microplate reader (Promega, GM3000).

### Western blot and immunoprecipitation

Immunoblotting and immunoprecipitation were performed according to standard protocols. Cells were treated with indicated compounds and then washed with cold PBS and lysed on ice for 5 min in NP40 buffer (50 mM Tris pH 7.5, 10% Glycerol, 1% NP40, 150 mM NaCl, 1 mM EDTA). For protein extraction from tumors obtained from *in vivo* studies, samples were washed with cold PBS and homogenized in RIPA buffer using the TissueRuptor II (Qiagen, 9002755). All lysis buffers were supplemented with protease inhibitors (Roche, 11836170001) and phosphatase inhibitors (Roche, 04906837001). Lysates were clarified by centrifugation at 15,000 rpm for 10 min at 4°C, and the protein concentration was quantified using the Pierce BCA protein assay kit (Thermo Fisher Scientific, 23227). Equal amounts of proteins were loaded and resolved on NuPAGE, 4–12% Bis-Tris polyacrylamide gels (Thermo Fisher Scientific, NP0335BOX), and transferred to nitrocellulose membranes (Cytiva, 10600006). After blocking the membranes in 5% milk, the membranes were incubated with primary antibodies overnight at 4°C. Antibodies against pMEK1/2^Ser217/221^ (#9154), MEK1 (#2352), pERK1/2^Thr202/Tyr204^ (#4370), ERK1/2 (#9102), ARAF (#75804), N/K ATPase (#3010), DUSP4 (#5149), DUSP6 (#50945), cyclin D1 (#55506), were from Cell Signaling Technology. Antibodies against BRAF (sc-5284) and HSP90 (sc-13119) were from Santa Cruz Biotechnology. BRAF(V600E) antibody was from NewEast Biosciences (26039). RAS was from Thermo Fisher Scientific (1862335). HA antibody was from Abcam (ab236632). KRAS antibody (NBP2-45536) was from Novus Biologicals. CRAF antibody was from BD Transduction Laboratories (610152), and pCRAF^Ser338^ antibody from Millipore Sigma (05-538). The next day, membranes were washed with TBS-T and incubated with anti-rabbit IgG HRP-linked (Cell Signaling Technology, #7074) or anti-mouse IgG HRP-linked (Cell Signaling Technology, #7076) secondary antibodies for 1 hour at room temperature. Chemiluminescent signals were detected on BioExcell® X-ray films (Worl Wide Medical Products, 41101001). In addition to MEK, β-Actin HRP-conjugated (Cell Signaling Technology, #5125) was used as a loading control. For immunoprecipitations, 1500 μg of proteins from lysates were incubated with the indicated anti-MEK1 (Cell Signaling Technology, #2352) or anti-BRAF (Santa Cruz Biotechnology, sc-5284) antibodies overnight at 4°C, followed by incubation with protein A (Thermo Fisher Scientific, 20333) or protein G agarose (Thermo Fisher Scientific, 20398) for 2 hours at 4°C. Samples were washed three times with cold lysis buffer, and sample buffer was added for subsequent immunoblotting analysis.

### RAS-GTP pull-down

Cells were treated with indicated compounds and RAS-GTP levels were estimated by pull-down assays using the Active Ras Pull-Down and Detection Kit (Thermo Fisher Scientific, 16117). Active RAS was analyzed by Western blot as previously described using the specific anti-RAS antibody provided with the kit.

### Cell fractionation

Cells were treated with compounds as indicated and cell fractionation experiments were performed using the ProteoExtract ® Subcellular Proteome extraction kit (Millipore Sigma, 539790). Cytosolic (fraction 1) and membrane (fraction 2) fractions were analyzed by Western blot.

### NanoBRET target engagement assay

BRAF-NanoLuc was obtained using QuikChange II XL-site directed mutagenesis kit (Agilent, 200522) and BRAF(V600E)-NanoLuc® fusion vector (Promega, NV2481) as a template. pMEV-2HA was purchased from Biomyx Technology (P1001), and full-length NRAS was cloned into the plasmid by restriction enzyme digestion. pMEV-NRAS(Q61R)-2HA was also obtained using the QuikChange II XL-site directed mutagenesis kit. 2 × 10^5^ 293-H cells, KRAS(Q61K) Tet-On MEFs, or SK-MEL-2 cells were co-transfected in suspension with 5 ng of BRAF-NanoLuc and 45 ng of pMEV-NRAS(WT)-2HA, pMEV-NRAS(Q61R)-2HA, or control transfection carrier DNA (Promega, E4881) using FuGENE HD transfection reagent (Promega, E2311), and were plated in white 96-well plates (Stellar Scientific, TC-CA-30196) with or without doxycycline. 24 hours later, cells were incubated with increasing concentrations of pan-RAFi-based tracer for 2 hours at 37°C. Where indicated, cells were treated with DMSO (control), cobimetinib, avutometinib, or daraxonrasib for 2 hours at 37°C before the addition of the pan-RAFi-based tracers. 20 μM fluorofurimazine (MedchemExpress, HY-D1282) was added to each well, and the plate was incubated for 2-3 minutes at room temperature before measuring donor emission wavelength (450 nm) and acceptor emission wavelength (610 nm) using the GloMax® Discover (Promega, GM3000) luminometer.

### NanoBiT protein-protein interaction assay

N-terminal fusions of small BiT (Sm-BiT)-KRAS and large BiT (Lg-BiT)-BRAF were cloned into the pMEV-2HA (Biomyx Technology, P1001) and pcDNA3, respectively. pcDNA3-LgBiT-BRAF(I666R) was obtained using the QuikChange II XL-site directed mutagenesis kit. 1 × 10^5^ 293-H cells were co-transfected in suspension with 25 ng of pcDNA3-LgBiT-BRAF or pcDNA3-LgBiT-BRAF(I666R) and 25 ng of pMEV-SmBiT-KRAS-2HA, using FuGENE HD transfection reagent (Promega, E2311), and were plated in white 96-well plates (Stellar Scientific, TC-CA-30196). 24 hours later, cells were treated with DMSO (control) or the indicated inhibitors, and luminescence signal was measured using the GloMax® Discover (Promega, GM3000) luminometer after adding 20 μM fluorofurimazine to each well.

### Animal xenograft models

All animals were examined before the initiation of studies to ensure that they were healthy and acclimated to the laboratory environment. 6-8-week-old female BALB/c nude mice (Shanghai BK Laboratory Animal Technology and Vital River Laboratories) or athymic Nude-*Foxn1^nu^* mice (Envigo Laboratories) were used for animal experiments. All mouse experiments were approved by the Institutional Animal Care and Use Committee (IACUC) of WuXi AppTec, Shanghai ChemPartner or the Icahn School of Medicine at Mount Sinai (protocol no. IACUC–2016–0066). Mice were maintained under specific pathogen-free conditions, and food and water were provided ad libitum. SK-MEL-2, NCI-H2122, or LS513 cells were maintained *in vitro* as a monolayer culture, and cells in the exponential growth phase were harvested, counted, and resuspended for tumor inoculation. Each mouse was inoculated subcutaneously in the right flank with 10 × 10^6^ SK-MEL-2 tumor cells, 10 × 10^6^ NCI-H2122 tumor cells, or 10 × 10^6^ LS513 tumor cells per injection. After inoculation, mice were monitored daily, weighed every three days, and caliper measurements began when tumors became visible. Tumor volume was calculated using the following formula: tumor volume = (D x d2)/2, where D and d refer to the long and short tumor diameters, respectively. When tumors reached a volume of 250 mm^³^, mice were randomized into 3 or 4 groups, depending on the experimental design for each xenograft model, to begin treatments as indicated. Avutometinib was prepared in 5% DMSO + 10% 2-hydroxypropyl-b-cyclodextrine (Millipore Sigma, H107) in Milli-Q water. Belvarafenib was prepared in 5% DMSO + 5% Cremophor® EL (Millipore Sigma, 238470) in Milli-Q water. Cobimetinib was prepared in 0.5% methyl cellulose (Millipore Sigma, M0512) + 0.2% Tween 80 (Millipore Sigma, 9490) in Milli-Q water. Mice were euthanized at the indicated time points, and tumors were collected and segmented for immunohistochemistry (IHQ) evaluation, bulk RNA sequencing, or Western blot. Endpoint criteria for tumor growth studies included tumor volume exceeding 1,000 mm^³^ or tumor ulceration. Results of mouse experiments are expressed as mean ± SEM of tumor volume for all tumors analyzed.

### Histopathology and immunohistochemical staining

Tumors were washed with cold PBS and placed in 10% formalin (Sigma-Aldrich, F5554) for 24 hours at room temperature and transferred to 70% ethanol. Subsequently, samples were paraffin embedded, sectioned (5 µm), and stained with hematoxylin and eosin (H&E) or anti-pERK1/2^Thr202/Tyr204^ (Cell Signaling Technology, #4370) according to standard protocols by the Histology and Immunostaining facility from the Department of Oncological Sciences at the Icahn School of Medicine at Mount Sinai.

### RNA sequencing library preparation

After washing the tumors with cold PBS, samples were stored in RNA*later* (Invitrogen, AM7020) prior to RNA extraction. Samples were homogenized in TRIzol reagent (Invitrogen, 15596026) using the TissueRuptor II, and RNA was isolated and treated with Turbo DNase (Invitrogen, AM1907) according to the manufacturers’ instructions. RNA quality was evaluated using the 2100 Bioanalyzer (Agilent, G2939A) to confirm suitability for poly(A)-selection-based library preparation. Libraries were then prepared with the NEBNext® Ultra II Directional RNA Library Prep Kit for Illumina (New England Biolabs, E7760S) using 500 ng of RNA input. Final libraries were sequenced on a NextSeq 2000 instrument using NextSeq 1000/2000 P2 Reagents (100 cycles) v3 (Illumina, 20046811), with 122 bp for read 1 and 8 bp dual-index sequencing.

### RNA sequencing data processing and analysis

Raw sequencing read quality was assessed using FastQC v0.11.9. Adapter trimming and removal of low-quality bases were performed using Trim Galore v0.6.6. Trimmed reads were aligned to the human genome GRCh38.p14 (GENCODE v46) using the STAR aligner v2.7.10a. The resulting transcriptome-aligned BAM files were used for transcript-level quantification, using Salmon v1.4.0 in alignment-based mode. Differential expression analysis was conducted using gene-level read counts and the DESeq2 v1.42.1 R package. Genes were considered differentially expressed if the adjusted *p*-value was < 0.05 and the absolute log2 fold change exceeded 1. Heatmaps of custom gene sets were generated using the pheatmap v1.0.12 R package.

### Statistical Analysis

Data are presented as mean ± SEM, and statistical comparisons were performed using a Mann-Whitney U test, a two-tailed unpaired Welch’s *t* test, or a one-way ANOVA test in GraphPad Prism 10, as indicated in the figure legends.

### Data availability

All requests for raw data and materials will be promptly reviewed by P.I. Poulikakos to determine if they are subject to intellectual property or confidentiality obligations. Any data and materials that can be shared will be released via a material transfer agreement.

## FIGURE LEGENDS

**Figure S1.**
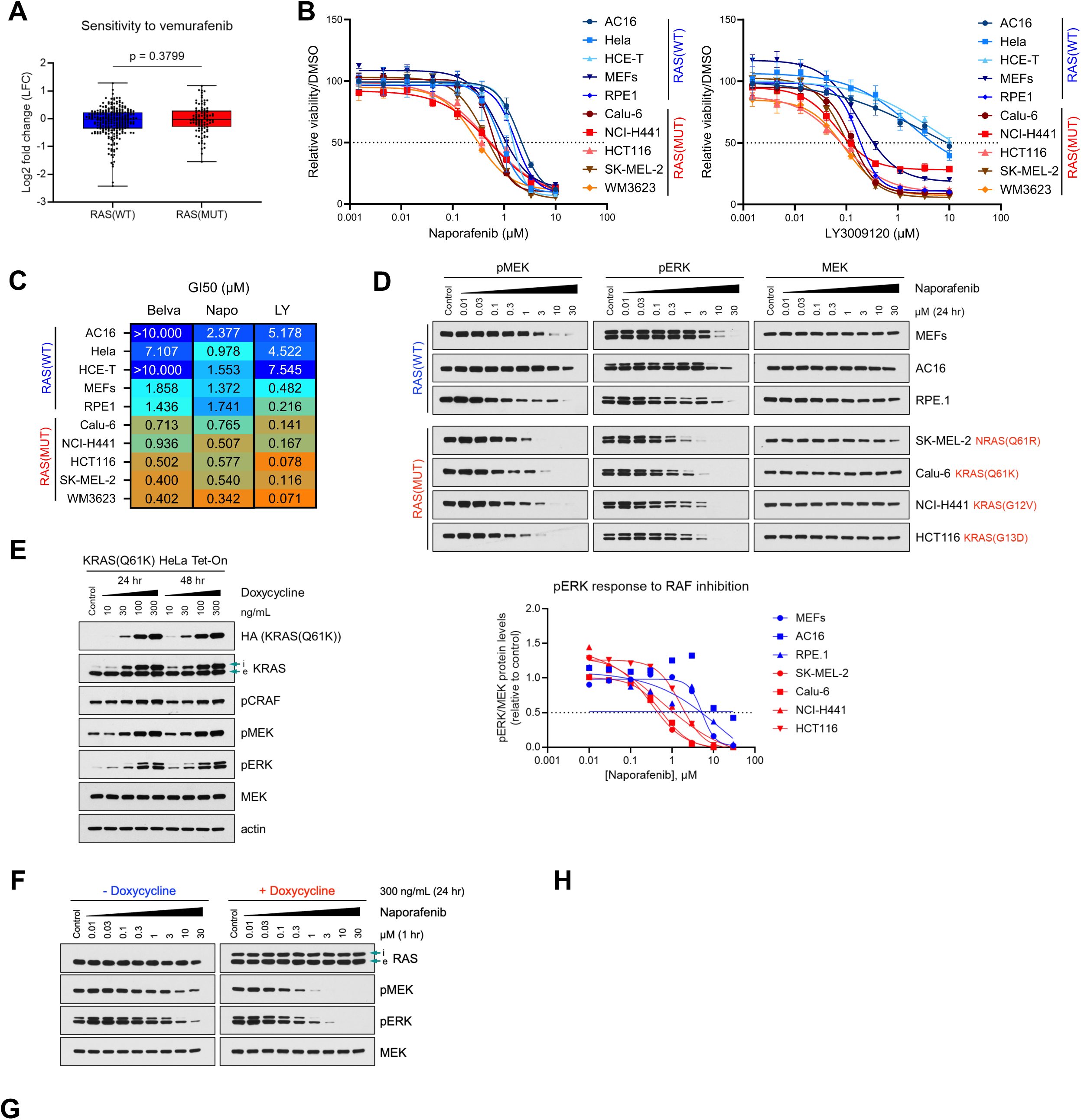
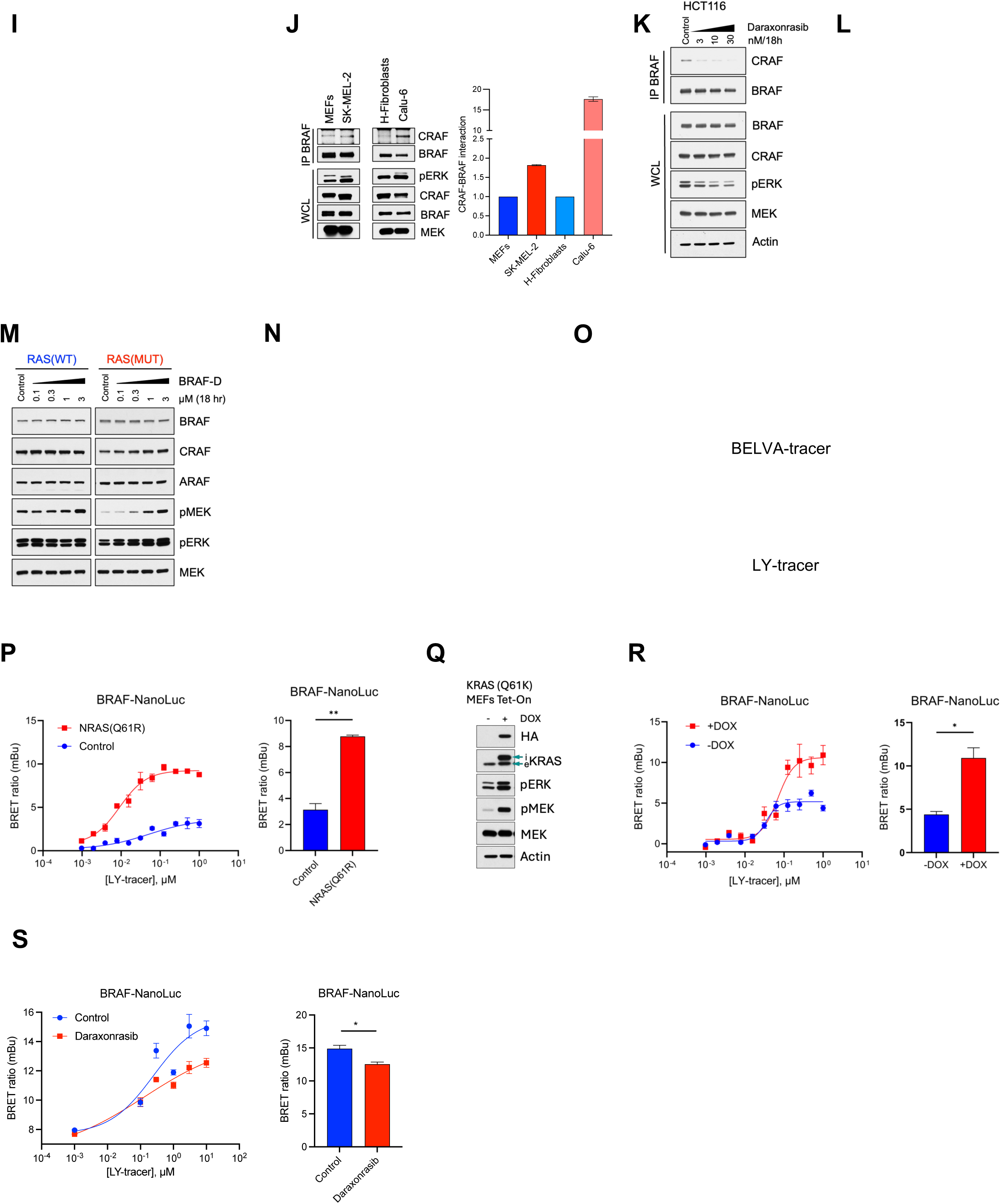
(related to Fig. 1) **A,** Box and whisker plot shows the distribution of drug response to the RAF inhibitor vemurafenib in RAS(WT) and RAS(MUT) cells from the PRISM Repurposing DepMap Public 24Q2 dataset. *P*-value was calculated using a Mann-Whitney U test in GraphPad Prism 10. **B,** Cell growth response to the pan-RAFis naporafenib and LY3009120 of the indicated RAS(WT) and RAS(MUT) cell lines after 6 days of treatment. Data are represented as mean ± SEM. **C,** Growth inhibition 50 (GI50) values in μM of belvarafenib, naporafenib, and LY3009120 estimated from dose-response curves by non-linear regression in the indicated cell lines using GraphPad Prism 10. **D,** The indicated RAS(WT) and RAS(MUT) cell lines were treated with DMSO (control) or increasing concentrations of naporafenib for 24 hours, and cell lysates were immunoblotted for pMEK, pERK, and MEK. The graph to the bottom shows the quantification of pERK normalized by the loading control MEK and relative to the control after ImageJ analysis. **E,** KRAS(Q61K) Tet-On HeLa cells were treated or not with increasing concentrations of doxycycline (DOX) for 24 or 48 hours, and cell lysates were immunoblotted for HA, KRAS (i, induced; e, endogenous), pCRAFS338, pMEK, pERK, MEK, and actin. **F,** KRAS(Q61K) Tet-On HeLa cells were treated or not with 300 ng/mL doxycycline (DOX) for 24 hours, followed by treatment with increasing concentrations of naporafenib for 1 hour, and cell lysates were immunoblotted for RAS (i, induced; e, endogenous), pMEK, pERK, and MEK. **G,** Chemical structure of pan-RAF-D. **H,** HCT116 and Calu-6 cells were treated with the indicated concentrations of VHL ligand, LY3009120 (LY1), bortezomib (BZ), or MLN4924 (MLN) for 2 hours, followed by treatment with 0.5 µM pan-RAF-D for 18 hours. Cell lysates were immunoblotted for BRAF, MEK, and actin. **I,** Panel of RAS(WT) and RAS(MUT) cells were treated with the indicated concentrations of pan-RAF-D for 18 hours. Cell lysates were immunoblotted for BRAF, CRAF, and MEK. **J**, Cell lysates from the indicated cell lines were either subjected to immunoprecipitation with a BRAF antibody followed by immunoblotting for CRAF and BRAF or immunoblotted with the indicated antibodies. Graph to the right shows the CRAF/BRAF ratio from IP BRAF. **K**, HCT116 cells were treated with increasing concentrations of daraxonrasib for 18 hours. Cell lysates were either subjected to immunoprecipitation with a BRAF antibody followed by immunoblotting for CRAF and BRAF or immunoblotted with the indicated antibodies. **L**, HCT116 cells were treated with increasing concentrations of the multi-RAS(ON)-inhibitor daraxonrasib alone or in combination with pan-RAF-D for 18 hours. Cell lysates were immunoblotted with the indicated antibodies. **M**, RAS(WT) and RAS(MUT) cells (MEFs and HCT116, respectively) were treated with DMSO (control) or increasing concentrations of BRAF-D for 18 hours. Cell lysates were immunoblotted for BRAF, CRAF, ARAF, pMEK, pERK, and MEK. **N,** A-375 cells were treated with DMSO (control) or increasing concentrations of pan-RAF-D or BRAF-D for 18 hours. Cell lysates were immunoblotted for BRAF(V600E), CRAF, ARAF, pMEK, pERK, and MEK. **O,** Chemical structures of the pan-RAFi-based tracers BELVA-tracer and LY-tracer. **P,** Dose-dependent BRET signal measured in 293-H cells co-transfected with BRAF-NanoLuc and pMEV-NRAS(Q61K)-2HA or a control transfection carrier DNA and incubated with LY-tracer. The graph on the right shows BRET signal at tracer concentration of 1 µM. Data are represented as mean ± SEM, and statistical comparison was performed using a two-tailed unpaired Welch’s *t* test in GraphPad Prism 10 (**, *P* < 0.01). **Q,** KRAS(Q61K) Tet-On MEFs were treated or not with 300 ng/mL doxycycline (DOX) for 24 hours, and cell lysates were immunoblotted for HA, KRAS (i, induced; e, endogenous), pMEK, pERK, MEK, and actin. **R**, Dose-dependent BRET signal measured in KRAS(Q61K) Tet-On MEFs transfected with BRAF-NanoLuc and treated or not with 300 ng/mL doxycycline (DOX) for 24 hours incubating them with LY-tracer. The graph on the right shows BRET signal at tracer concentration of 1 µM. Data are represented as mean ± SEM, and statistical comparison was performed using a two-tailed unpaired Welch’s *t* test in GraphPad Prism 10 (*, *P* < 0.05). **S**, Dose-dependent BRET signal measured in SK-MEL-2 cells transfected with BRAF-NanoLuc, treated with DMSO (control) or with 10 nM daraxonrasib for 2 hours and incubated with LY-tracer. The graph on the right shows BRET signal at tracer concentration of 10 µM. Data are represented as mean ± SEM, and statistical comparison was performed using a two-tailed unpaired Welch’s *t* test in GraphPad Prism 10 (*, *P* < 0.05).

**Figure S2.**
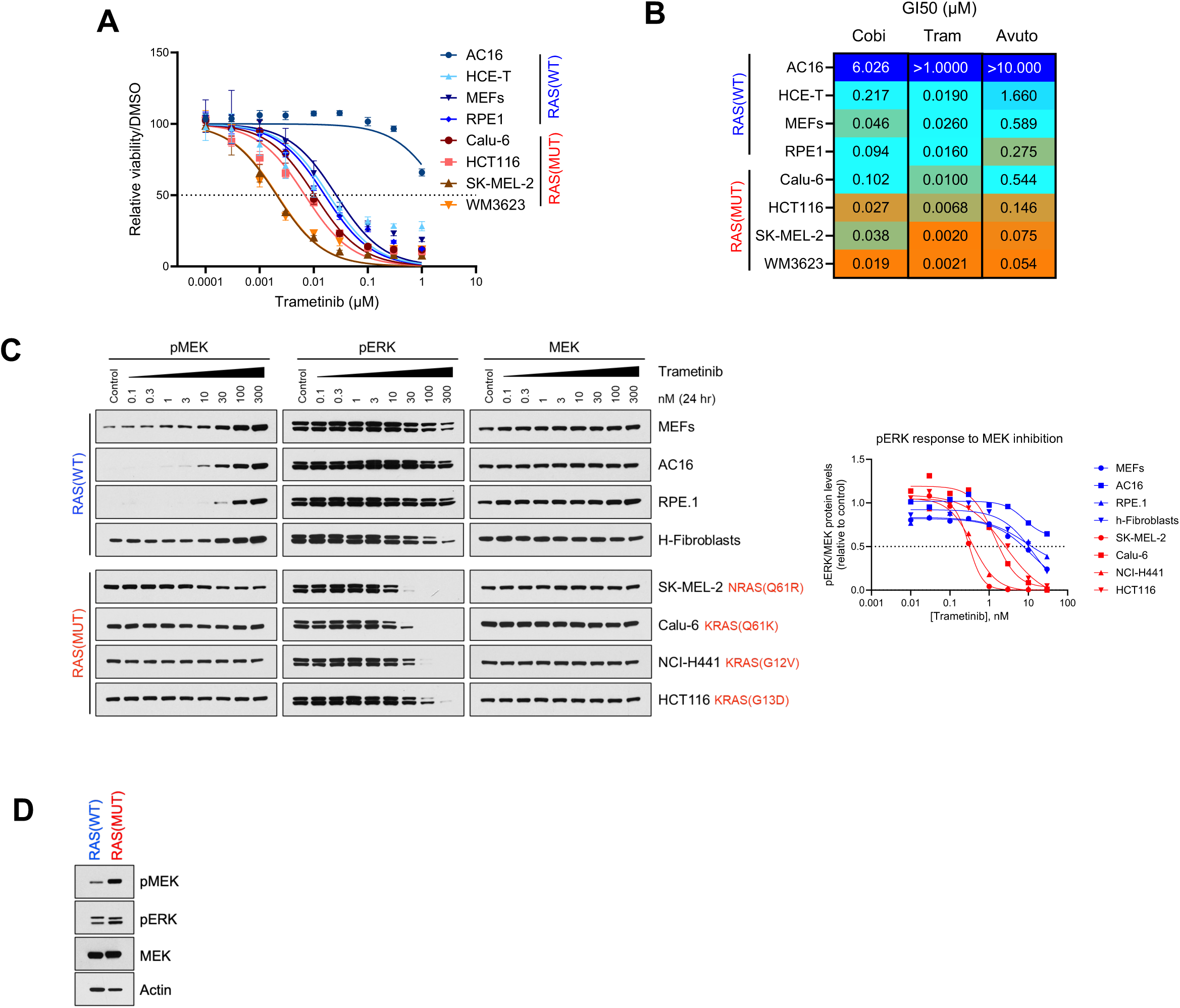
(related to Fig. 2) **A,** Cell growth response to trametinib of the indicated RAS(WT) and RAS(MUT) cell lines after 6 days of treatment. Data are represented as mean ± SEM. **B,** Growth inhibition 50 (GI50) values in μM of cobimetinib, trametinib, and avutometinib estimated from dose-response curves by non-linear regression in the indicated cell lines using GraphPad Prism 10. **C,** The indicated RAS(WT) and RAS(MUT) cell lines were treated with DMSO (control) or increasing concentrations of trametinib for 24 hours, and cell lysates were immunoblotted for pMEK, pERK, and MEK. The graph on the right shows the quantification of pERK normalized by the loading control MEK and relative to the control after ImageJ analysis. **D,** Cell lysates from RAS(WT) and RAS(MUT) cells (MEFs and HCT116, respectively) were immunoblotted for basal level comparison of pMEK, pERK, MEK, and actin.

**Figure S3.**
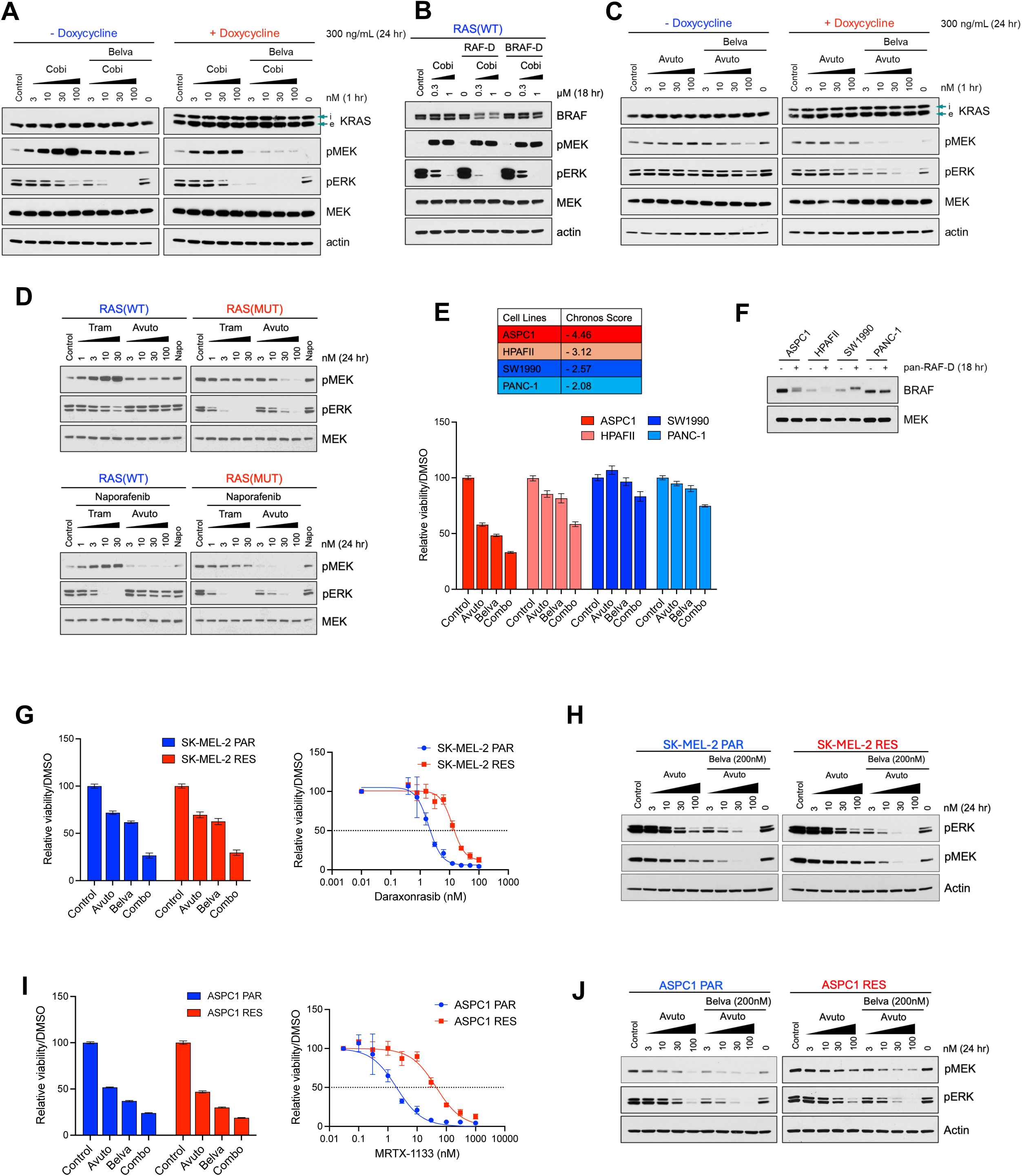
(related to Fig. 3) **A,** KRAS(Q61K) Tet-On MEFs were treated with or without 300 ng/mL doxycycline for 24 hours followed by treatment with DMSO (control), 200 nM belvarafenib (belva), or increasing concentrations of cobimetinib (cobi) alone or in combination with 200 nM belvarafenib for 1 hour, and cell lysates were immunoblotted for KRAS (i, induced; e, endogenous), pMEK, pERK, MEK, and actin. **B,** MEFs were treated with DMSO (control) or increasing concentrations of cobimetinib (cobi) alone or in combination with 0.5 µM pan-RAF-D or 0.5 µM BRAF-D for 18 hours, and cell lysates were immunoblotted for BRAF, pMEK, pERK, MEK, and actin. **C,** KRAS(Q61K) Tet-On MEFs were treated with or without 300 ng/mL doxycycline for 24 hours followed by treatment with DMSO (control), 200 nM belvarafenib (belva), or increasing concentrations of avutometinib (avuto) alone or in combination with 200 nM belvarafenib for 1 hour, and cell lysates were immunoblotted for KRAS (i, induced; e, endogenous), pMEK, pERK, MEK, and actin. **D,** RAS(WT) and RAS(MUT) cells (MEFs and SK-MEL-2 cells, respectively) were treated with DMSO (control), 200 nM naporafenib (napo), increasing concentrations of trametinib (tram) alone or in combination with 200 nM naporafenib, or increasing concentrations of avutometinib (avuto) alone or in combination with 200 nM naporafenib for 24 hours. Cell lysates were immunoblotted for pMEK, pERK, and MEK. **E**, Table on the top shows KRAS chronos scores from the DepMap portal for the indicated RAS-dependent (ASPC1 and HPAFII) and RAS-independent (SW1990 and PANC-1) cell lines. The graph on the bottom shows cell growth response to 250 nM belvarafenib (belva), 30 nM avutometinib (avuto), or the combination in the indicated cell lines after 6 days of treatment. Data are represented as mean ± SEM. **F**, The indicated cell lines were treated with 1µM pan-RAF-D for 18 hours. Cell lysates were immunoblotted with BRAF and MEK. **G**, Cell growth response to 250 nM belvarafenib (belva), 30 nM avutometinib (avuto), or the combination in SK-MEL-2 parental cells and derived cells resistant to daraxonrasib after 6 days of treatment. The graph on the right shows cell growth response to daraxonrasib in SK-MEL-2 parental and daraxonrasib resistant cells after 6 days of treatment. Data are represented as mean ± SEM. **H**, SK-MEL-2 parental and daraxonrasib resistant cells were treated with DMSO (control), 200 nM belvarafenib (belva), or increasing concentrations of avutometinib (avuto) alone or in combination with 200 nM belvarafenib for 24 hours, and cell lysates were immunoblotted for pMEK, pERK, and actin. **I**, Cell growth response to 250 nM belvarafenib (belva), 30 nM avutometinib (avuto), or the combination in ASPC1 parental cells and derived cells resistant to MRTX-1133 after 6 days of treatment. The graph on the right shows cell growth response to MRTX-1133 in ASPC1 parental and MRTX-1133 resistant cells after 6 days of treatment. Data are represented as mean ± SEM. **J**, ASPC1 parental and MRTX-1133 resistant cells were treated with DMSO (control), 200 nM belvarafenib (belva), or increasing concentrations of avutometinib (avuto) alone or in combination with 200 nM belvarafenib for 24 hours, and cell lysates were immunoblotted for pMEK, pERK, and actin.

**Figure S4.**
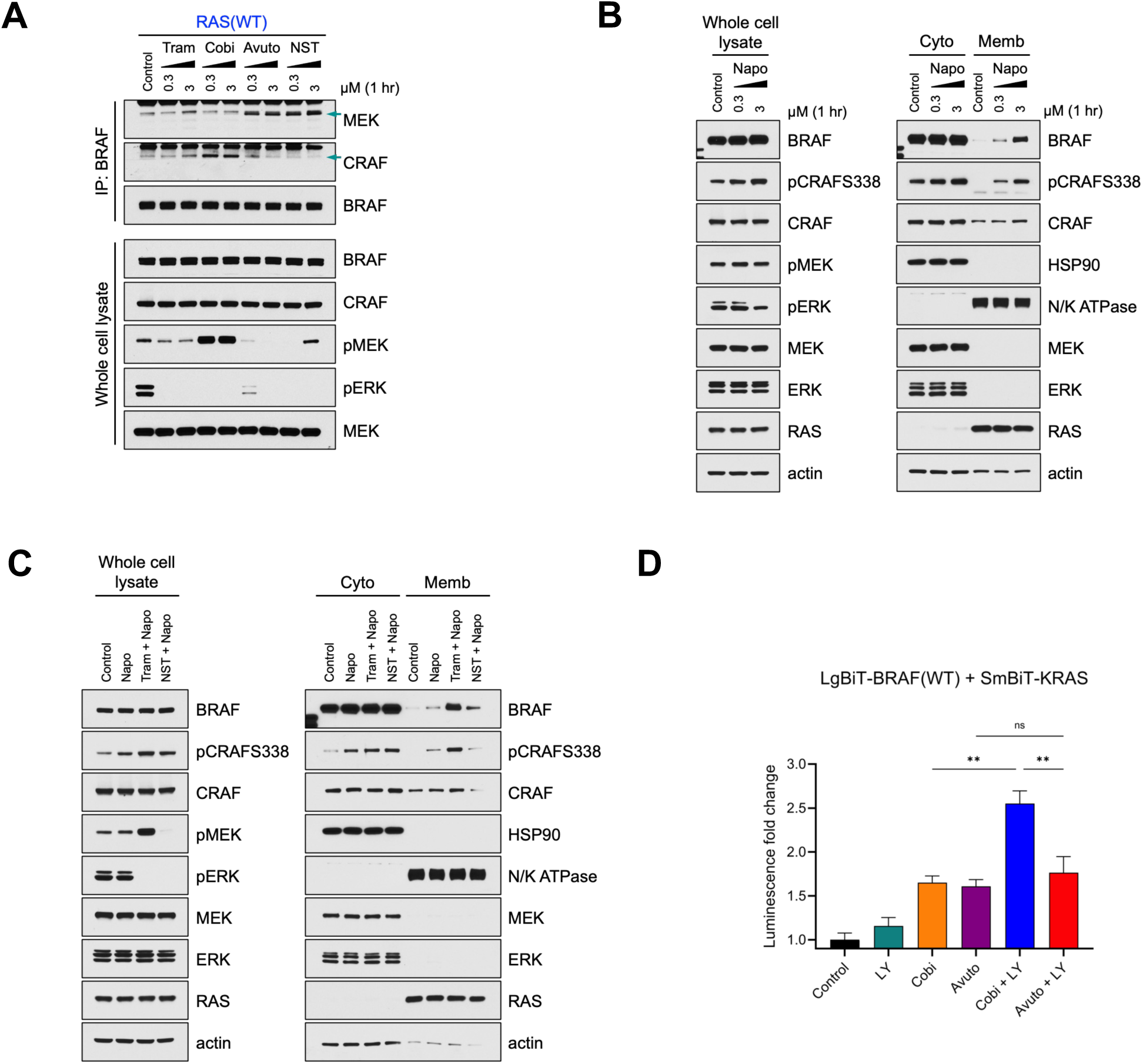
(related to Fig. 4) **A,** MEFs were treated with increasing concentrations of trametinib (tram), cobimetinib (cobi), avutometinib (avuto), or NST-628 (NST) for 1 hour. Cell lysates were subjected to immunoprecipitation with a BRAF antibody, followed by immunoblotting for MEK, CRAF, and BRAF, or were immunoblotted with the indicated antibodies. **B,** HeLa cells were treated with increasing concentrations of naporafenib (napo) for 1 hour. Cells underwent cell fractionation, and cytosolic (cyto) and membrane (memb) fractions, along with whole cell lysates, were immunoblotted with the indicated antibodies. **C,** HeLa cells were treated with 0.3 µM naporafenib (napo) for 1 hour or with 0.3 µM trametinib (tram) or 0.3 µM NST-628 (NST) for 1 hour, followed by treatment with 0.3 µM naporafenib for 1 hour. Cells underwent cell fractionation, and cytosolic (cyto) and membrane (memb) fractions, along with whole cell lysates, were immunoblotted with the indicated antibodies. **D,** Luminescence signal fold change measured in 293-H cells co-transfected with LgBiT-BRAF and SmBiT-KRAS, and treated with 0.3 µM LY3009120 (LY) for 1 hour, 3 µM cobimetinib (cobi) for 2 hours, 3 µM avutometinib (avuto) for 2 hours, or with 3 µM cobimetinib or 3 µM avutometinib for 1 hour, followed by treatment with 0.3 µM LY3009120 for 1 hour. Data are represented as mean ± SEM. Statistical comparison was performed using a one-way ANOVA test in GraphPad Prism 10 (ns, non-significant; *, *P* < 0.05).

**Figure S5.**
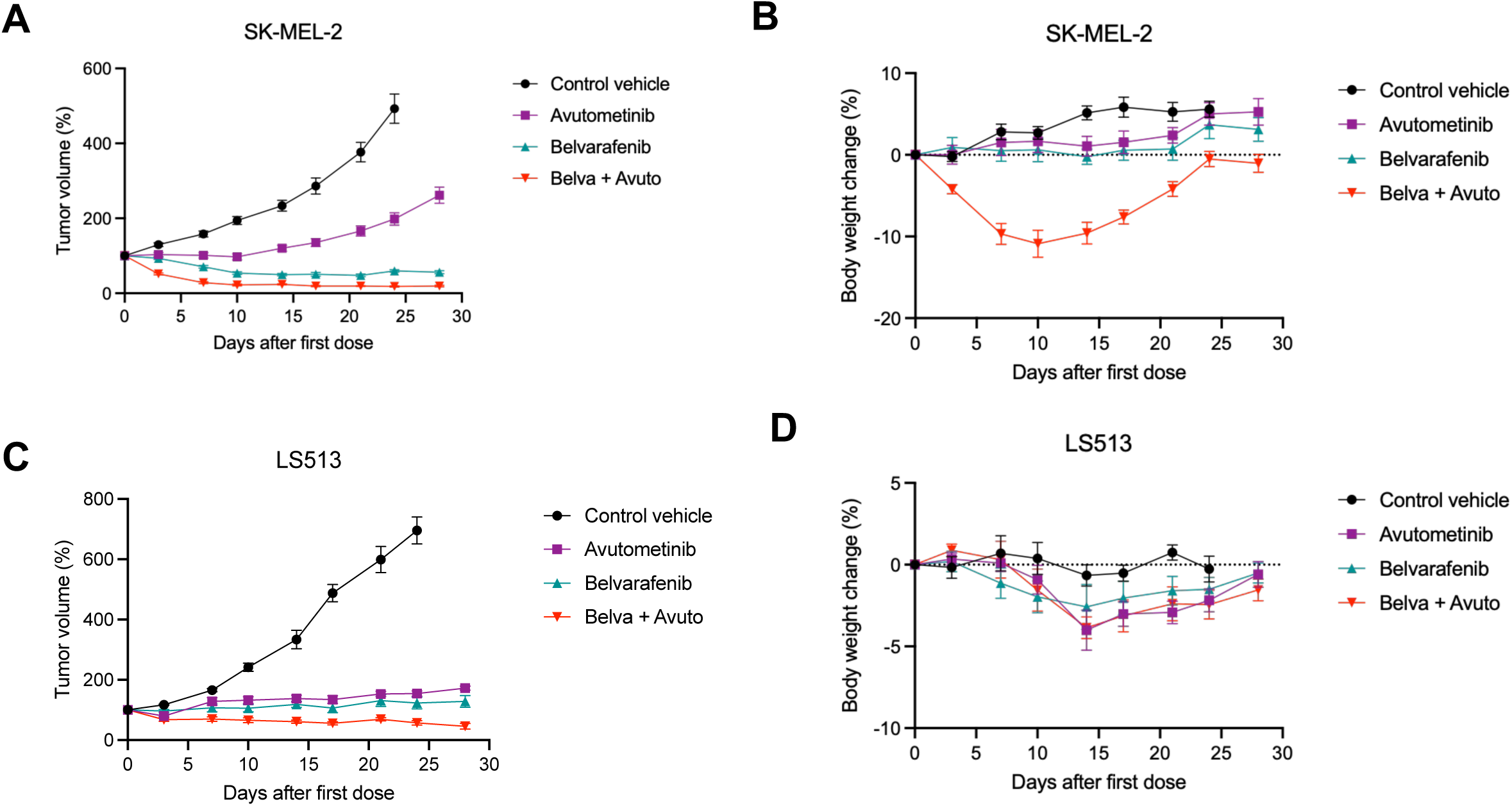
(related to Fig. 5) **A,** SK-MEL-2 xenograft growth kinetics in female BALB/c nude mice treated with (i) control vehicle, (ii) avutometinib 0.1 mg/kg for 5 days on (QDx5), 2 days off, administered orally (PO), (iii) belvarafenib 15 mg/kg once daily (QD) PO, or (iv) belvarafenib 15 mg/kg QD PO combined with avutometinib 0.1 mg/kg QDx5, 2 days off, PO for 28 days. Data are presented as mean ± SEM, n ≥ 8 mice/group. **B,** The change in body weight from **A**. Data are presented as mean ± SEM, n ≥ 8 mice/group. **C,** LS513 xenograft kinetics in female BALB/c nude mice treated as in **A**. Data are presented as mean ± SEM, n = 8 mice/group. **D,** The change in body weight from **C**. Data are presented as mean ± SEM, n = 8 mice/group.

**Figure S6.**
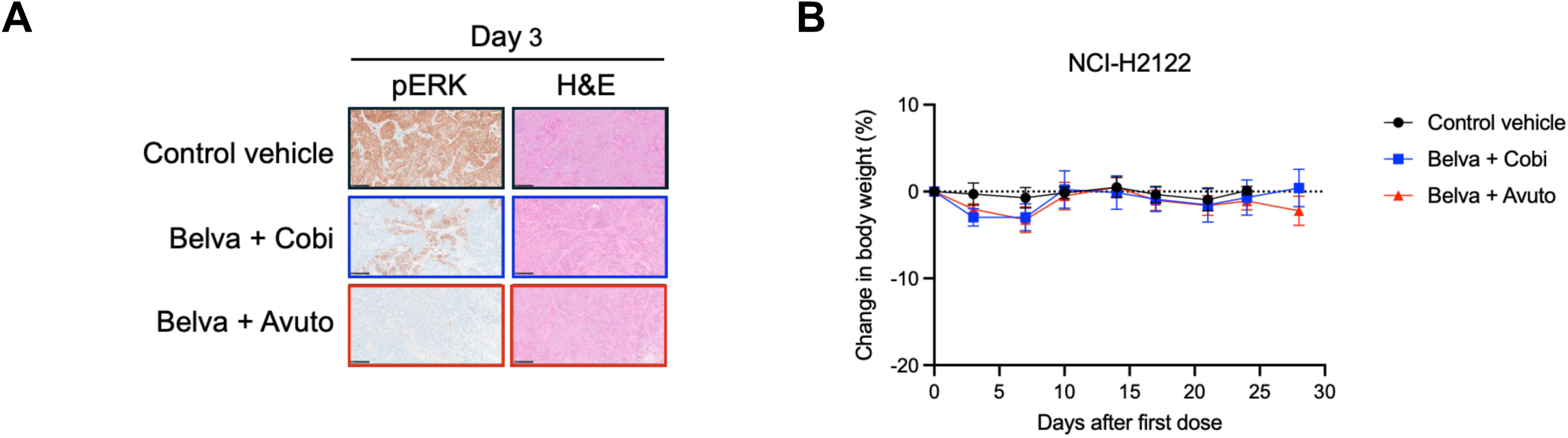
(related to Fig. 6) **A,** Representative immunohistochemistry (IHC) against pERK and hematoxylin and eosin (H&E) staining of tumor sections from female athymic Nude-*Foxn1^nu^* mice bearing NCI-H2122 xenografts and treated with (i) control vehicle, (ii) belvarafenib 15 mg/kg once daily (QD), administered orally (PO) combined with cobimetinib 5 mg/kg QD PO, or (iii) belvarafenib 15 mg/kg QD PO combined with avutometinib 0.3 mg/kg QD PO, for 3 days. **B,** The change in body weight of female athymic Nude-*Foxn1^nu^* mice bearing NCI-H2122 xenografts and treated as in **B** for 28 days. Data are presented as mean ± SEM, n = 5 mice/ control vehicle group, n = 8 mice/belva + cobi group, n = 9 mice/belva + avuto group.

## References

1. Matallanas D, Birtwistle M, Romano D, Zebisch A, Rauch J, von Kriegsheim A, et al. Raf family kinases: old dogs have learned new tricks. Genes & cancer 2011;2(3):232–60 doi 10.1177/1947601911407323.

2. Baines AT, Xu D, Der CJ. Inhibition of Ras for cancer treatment: the search continues. Future medicinal chemistry 2011;3(14):1787–808 doi 10.4155/fmc.11.121.

3. Poulikakos PI, Sullivan RJ, Yaeger R. Molecular Pathways and Mechanisms of BRAF in Cancer Therapy. Clinical cancer research : an official journal of the American Association for Cancer Research 2022 doi 10.1158/1078-0432.CCR-21-2138.

4. Samatar AA, Poulikakos PI. Targeting RAS-ERK signalling in cancer: promises and challenges. Nature reviews Drug discovery 2014;13(12):928–42 doi 10.1038/nrd4281.

5. Young A, Lyons J, Miller AL, Phan VT, Alarcon IR, McCormick F. Ras signaling and therapies. Advances in cancer research 2009;102:1–17 doi 10.1016/S0065-230X(09)02001-6.

6. Falchook GS, Long GV, Kurzrock R, Kim KB, Arkenau TH, Brown MP, et al. Dabrafenib in patients with melanoma, untreated brain metastases, and other solid tumours: a phase 1 dose-escalation trial. Lancet 2012;379(9829):1893–901 doi 10.1016/S0140-6736(12)60398-5.

7. Chapman PB, Hauschild A, Robert C, Haanen JB, Ascierto P, Larkin J, et al. Improved survival with vemurafenib in melanoma with BRAF V600E mutation. The New England journal of medicine 2011;364(26):2507–16 doi 10.1056/NEJMoa1103782.

8. Holderfield M, Deuker MM, McCormick F, McMahon M. Targeting RAF kinases for cancer therapy: BRAF-mutated melanoma and beyond. Nature reviews Cancer 2014;14(7):455–67 doi 10.1038/nrc3760.

9. Karoulia Z, Gavathiotis E, Poulikakos PI. New perspectives for targeting RAF kinase in human cancer. Nature reviews Cancer 2017;17(11):676–91 doi 10.1038/nrc.2017.79.

10. Larkin J, Ascierto PA, Dreno B, Atkinson V, Liszkay G, Maio M, et al. Combined vemurafenib and cobimetinib in BRAF-mutated melanoma. The New England journal of medicine 2014;371(20):1867–76 doi 10.1056/NEJMoa1408868.

11. Robert C, Karaszewska B, Schachter J, Rutkowski P, Mackiewicz A, Stroiakovski D, et al. Improved overall survival in melanoma with combined dabrafenib and trametinib. The New England journal of medicine 2015;372(1):30–9 doi 10.1056/NEJMoa1412690.

12. Dummer R, Ascierto PA, Gogas HJ, Arance A, Mandala M, Liszkay G, et al. Encorafenib plus binimetinib versus vemurafenib or encorafenib in patients with BRAF-mutant melanoma (COLUMBUS): a multicentre, open-label, randomised phase 3 trial. The lancet oncology 2018;19(5):603–15 doi 10.1016/S1470-2045(18)30142-6.

13. Poulikakos PI, Zhang C, Bollag G, Shokat KM, Rosen N. RAF inhibitors transactivate RAF dimers and ERK signalling in cells with wild-type BRAF. Nature 2010;464(7287):427–30 doi 10.1038/nature08902.

14. Poulikakos PI, Persaud Y, Janakiraman M, Kong X, Ng C, Moriceau G, et al. RAF inhibitor resistance is mediated by dimerization of aberrantly spliced BRAF(V600E). Nature 2011;480(7377):387–90 doi 10.1038/nature10662.

15. Karoulia Z, Wu Y, Ahmed TA, Xin Q, Bollard J, Krepler C, et al. An Integrated Model of RAF Inhibitor Action Predicts Inhibitor Activity against Oncogenic BRAF Signaling. Cancer cell 2016;30(3):485–98 doi 10.1016/j.ccell.2016.06.024.

16. Adamopoulos C, Ahmed TA, Tucker MR, Ung PMU, Xiao M, Karoulia Z, et al. Exploiting Allosteric Properties of RAF and MEK Inhibitors to Target Therapy-Resistant Tumors Driven by Oncogenic BRAF Signaling. Cancer discovery 2021;11(7):1716–35 doi 10.1158/2159-8290.CD-20-1351.

17. Monaco KA, Delach S, Yuan J, Mishina Y, Fordjour P, Labrot E, et al. LXH254, a Potent and Selective ARAF-Sparing Inhibitor of BRAF and CRAF for the Treatment of MAPK-Driven Tumors. Clinical cancer research : an official journal of the American Association for Cancer Research 2021;27(7):2061–73 doi 10.1158/1078-0432.CCR-20-2563.

18. Peng SB, Henry JR, Kaufman MD, Lu WP, Smith BD, Vogeti S, et al. Inhibition of RAF Isoforms and Active Dimers by LY3009120 Leads to Anti-tumor Activities in RAS or BRAF Mutant Cancers. Cancer cell 2015;28(3):384–98 doi 10.1016/j.ccell.2015.08.002.

19. Tang Z, Yuan X, Du R, Cheung SH, Zhang G, Wei J, et al. BGB-283, a Novel RAF Kinase and EGFR Inhibitor, Displays Potent Antitumor Activity in BRAF-Mutated Colorectal Cancers. Molecular cancer therapeutics 2015;14(10):2187–97 doi 10.1158/1535-7163.MCT-15-0262.

20. Chen SH, Zhang Y, Van Horn RD, Yin T, Buchanan S, Yadav V, et al. Oncogenic BRAF Deletions That Function as Homodimers and Are Sensitive to Inhibition by RAF Dimer Inhibitor LY3009120. Cancer discovery 2016;6(3):300–15 doi 10.1158/2159-8290.CD-15-0896.

21. Yen I, Shanahan F, Lee J, Hong YS, Shin SJ, Moore AR, et al. ARAF mutations confer resistance to the RAF inhibitor belvarafenib in melanoma. Nature 2021;594(7863):418–23 doi 10.1038/s41586-021-03515-1.

22. de Braud F, Dooms C, Heist RS, Lebbe C, Wermke M, Gazzah A, et al. Initial Evidence for the Efficacy of Naporafenib in Combination With Trametinib in NRAS-Mutant Melanoma: Results From the Expansion Arm of a Phase Ib, Open-Label Study. Journal of clinical oncology : official journal of the American Society of Clinical Oncology 2023;41(14):2651–60 doi 10.1200/JCO.22.02018.

23. Cassier P, Mehnert JM, Kim R, Chmielowski B, Millward M, Perez CA, et al. 613MO A phase I clinical trial evaluating exarafenib, a selective pan-RAF inhibitor in combination with binimetinib in NRAS-mutant (NRASMut) melanoma (Mel) & BRAF-altered solid tumors. Annals of Oncology 2024;35:S491–S2 doi 10.1016/j.annonc.2024.08.680.

24. A phase Ib trial of belvarafenib in combination with cobimetinib in patients with advanced solid tumors: Interim results of dose-escalation and patients with NRAS-mutant melanoma of dose-expansion. https://ascopubs.org/doi/10.1200/JCO.2021.39.15_suppl.3007.

25. Tsherniak A, Vazquez F, Montgomery PG, Weir BA, Kryukov G, Cowley GS, et al. Defining a Cancer Dependency Map. Cell 2017;170(3):564–76 e16 doi 10.1016/j.cell.2017.06.010.

26. Joseph EW, Pratilas CA, Poulikakos PI, Tadi M, Wang W, Taylor BS, et al. The RAF inhibitor PLX4032 inhibits ERK signaling and tumor cell proliferation in a V600E BRAF-selective manner. Proceedings of the National Academy of Sciences of the United States of America 2010;107(33):14903–8 doi 10.1073/pnas.1008990107.

27. Hatzivassiliou G, Song K, Yen I, Brandhuber BJ, Anderson DJ, Alvarado R, et al. RAF inhibitors prime wild-type RAF to activate the MAPK pathway and enhance growth. Nature 2010;464(7287):431–5 doi 10.1038/nature08833.

28. Halaban R, Zhang W, Bacchiocchi A, Cheng E, Parisi F, Ariyan S, et al. PLX4032, a selective BRAF(V600E) kinase inhibitor, activates the ERK pathway and enhances cell migration and proliferation of BRAF melanoma cells. Pigment cell & melanoma research 2010;23(2):190–200 doi 10.1111/j.1755-148X.2010.00685.x.

29. Ramurthy S, Taft BR, Aversa RJ, Barsanti PA, Burger MT, Lou Y, et al. Design and Discovery of N-(3-(2-(2-Hydroxyethoxy)-6-morpholinopyridin-4-yl)-4-methylphenyl)-2-(trifluoromethyl)isonicotinamide, a Selective, Efficacious, and Well-Tolerated RAF Inhibitor Targeting RAS Mutant Cancers: The Path to the Clinic. Journal of medicinal chemistry 2020;63(5):2013–27 doi 10.1021/acs.jmedchem.9b00161.

30. Henry JR, Kaufman MD, Peng SB, Ahn YM, Caldwell TM, Vogeti L, et al. Discovery of 1-(3,3-dimethylbutyl)-3-(2-fluoro-4-methyl-5-(7-methyl-2-(methylamino)pyrido[2,3-d]pyrimidin-6-yl)phenyl)urea (LY3009120) as a pan-RAF inhibitor with minimal paradoxical activation and activity against BRAF or RAS mutant tumor cells. Journal of medicinal chemistry 2015;58(10):4165–79 doi 10.1021/acs.jmedchem.5b00067.

31. Bekes M, Langley DR, Crews CM. PROTAC targeted protein degraders: the past is prologue. Nature reviews Drug discovery 2022;21(3):181–200 doi 10.1038/s41573-021-00371-6.

32. Cregg J, Edwards AV, Chang S, Lee BJ, Knox JE, Tomlinson ACA, et al. Discovery of Daraxonrasib (RMC-6236), a Potent and Orally Bioavailable RAS(ON) Multi-selective, Noncovalent Tri-complex Inhibitor for the Treatment of Patients with Multiple RAS-Addicted Cancers. Journal of medicinal chemistry 2025;68(6):6064–83 doi 10.1021/acs.jmedchem.4c02314.

33. Holderfield M, Lee BJ, Jiang J, Tomlinson A, Seamon KJ, Mira A, et al. Concurrent inhibition of oncogenic and wild-type RAS-GTP for cancer therapy. Nature 2024 doi 10.1038/s41586-024-07205-6.

34. Alabi S, Jaime-Figueroa S, Yao Z, Gao Y, Hines J, Samarasinghe KTG, et al. Mutant-selective degradation by BRAF-targeting PROTACs. Nature communications 2021;12(1):920 doi 10.1038/s41467-021-21159-7.

35. Freeman AK, Ritt DA, Morrison DK. Effects of Raf dimerization and its inhibition on normal and disease-associated Raf signaling. Molecular cell 2013;49(4):751–8 doi 10.1016/j.molcel.2012.12.018.

36. Abe H, Kikuchi S, Hayakawa K, Iida T, Nagahashi N, Maeda K, et al. Discovery of a Highly Potent and Selective MEK Inhibitor: GSK1120212 (JTP-74057 DMSO Solvate). ACS medicinal chemistry letters 2011;2(4):320–4 doi 10.1021/ml200004g.

37. Rice KD, Aay N, Anand NK, Blazey CM, Bowles OJ, Bussenius J, et al. Novel Carboxamide-Based Allosteric MEK Inhibitors: Discovery and Optimization Efforts toward XL518 (GDC-0973). ACS medicinal chemistry letters 2012;3(5):416–21 doi 10.1021/ml300049d.

38. Hu J, Wei J, Yim H, Wang L, Xie L, Jin MS, et al. Potent and Selective Mitogen-Activated Protein Kinase Kinase 1/2 (MEK1/2) Heterobifunctional Small-molecule Degraders. Journal of medicinal chemistry 2020;63(24):15883–905 doi 10.1021/acs.jmedchem.0c01609.

39. Ishii N, Harada N, Joseph EW, Ohara K, Miura T, Sakamoto H, et al. Enhanced inhibition of ERK signaling by a novel allosteric MEK inhibitor, CH5126766, that suppresses feedback reactivation of RAF activity. Cancer research 2013;73(13):4050–60 doi 10.1158/0008-5472.CAN-12-3937.

40. Banerjee SN, Van Nieuwenhuysen E, Aghajanian C, D’Hondt V, Monk BJ, Clamp A, et al. Efficacy and Safety of Avutometinib +/- Defactinib in Recurrent Low-Grade Serous Ovarian Cancer: Primary Analysis of ENGOT-OV60/GOG-3052/RAMP 201. Journal of clinical oncology : official journal of the American Society of Clinical Oncology 2025;43(25):2782–92 doi 10.1200/JCO-25-00112.

41. Lito P, Saborowski A, Yue J, Solomon M, Joseph E, Gadal S, et al. Disruption of CRAF-mediated MEK activation is required for effective MEK inhibition in KRAS mutant tumors. Cancer cell 2014;25(5):697–710 doi 10.1016/j.ccr.2014.03.011.

42. Hatzivassiliou G, Haling JR, Chen H, Song K, Price S, Heald R, et al. Mechanism of MEK inhibition determines efficacy in mutant KRAS-versus BRAF-driven cancers. Nature 2013;501(7466):232–6 doi 10.1038/nature12441.

43. Khan ZM, Real AM, Marsiglia WM, Chow A, Duffy ME, Yerabolu JR, et al. Structural basis for the action of the drug trametinib at KSR-bound MEK. Nature 2020;588(7838):509–14 doi 10.1038/s41586-020-2760-4.

44. Lito P, Pratilas CA, Joseph EW, Tadi M, Halilovic E, Zubrowski M, et al. Relief of profound feedback inhibition of mitogenic signaling by RAF inhibitors attenuates their activity in BRAFV600E melanomas. Cancer cell 2012;22(5):668–82 doi 10.1016/j.ccr.2012.10.009.

45. Prahallad A, Sun C, Huang S, Di Nicolantonio F, Salazar R, Zecchin D, et al. Unresponsiveness of colon cancer to BRAF(V600E) inhibition through feedback activation of EGFR. Nature 2012;483(7387):100–3 doi 10.1038/nature10868.

46. Corcoran RB, Ebi H, Turke AB, Coffee EM, Nishino M, Cogdill AP, et al. EGFR-mediated re-activation of MAPK signaling contributes to insensitivity of BRAF mutant colorectal cancers to RAF inhibition with vemurafenib. Cancer discovery 2012;2(3):227–35 doi 10.1158/2159-8290.CD-11-0341.

47. Duncan JS, Whittle MC, Nakamura K, Abell AN, Midland AA, Zawistowski JS, et al. Dynamic reprogramming of the kinome in response to targeted MEK inhibition in triple-negative breast cancer. Cell 2012;149(2):307–21 doi 10.1016/j.cell.2012.02.053.

48. Wang X, Allen S, Blake JF, Bowcut V, Briere DM, Calinisan A, et al. Identification of MRTX1133, a Noncovalent, Potent, and Selective KRAS(G12D) Inhibitor. Journal of medicinal chemistry 2022;65(4):3123–33 doi 10.1021/acs.jmedchem.1c01688.

49. Park E, Rawson S, Li K, Kim BW, Ficarro SB, Pino GG, et al. Architecture of autoinhibited and active BRAF-MEK1-14-3-3 complexes. Nature 2019;575(7783):545–50 doi 10.1038/s41586-019-1660-y.

50. Martinez Fiesco JA, Durrant DE, Morrison DK, Zhang P. Structural insights into the BRAF monomer-to-dimer transition mediated by RAS binding. Nature communications 2022;13(1):486 doi 10.1038/s41467-022-28084-3.

51. Ryan MB, Quade B, Schenk N, Fang Z, Zingg M, Cohen SE, et al. The Pan-RAF-MEK Nondegrading Molecular Glue NST-628 Is a Potent and Brain-Penetrant Inhibitor of the RAS-MAPK Pathway with Activity across Diverse RAS- and RAF-Driven Cancers. Cancer discovery 2024;14(7):1190–205 doi 10.1158/2159-8290.CD-24-0139.

52. Leevers SJ, Paterson HF, Marshall CJ. Requirement for Ras in Raf activation is overcome by targeting Raf to the plasma membrane. Nature 1994;369(6479):411–4 doi 10.1038/369411a0.

53. Rajakulendran T, Sahmi M, Lefrancois M, Sicheri F, Therrien M. A dimerization-dependent mechanism drives RAF catalytic activation. Nature 2009;461(7263):542–5 doi 10.1038/nature08314.

54. Lavoie H, Sahmi M, Maisonneuve P, Marullo SA, Thevakumaran N, Jin T, et al. MEK drives BRAF activation through allosteric control of KSR proteins. Nature 2018;554(7693):549–53 doi 10.1038/nature25478.

55. Pratilas CA, Taylor BS, Ye Q, Viale A, Sander C, Solit DB, et al. (V600E)BRAF is associated with disabled feedback inhibition of RAF-MEK signaling and elevated transcriptional output of the pathway. Proceedings of the National Academy of Sciences of the United States of America 2009;106(11):4519–24 doi 10.1073/pnas.0900780106.

56. Dry JR, Pavey S, Pratilas CA, Harbron C, Runswick S, Hodgson D, et al. Transcriptional pathway signatures predict MEK addiction and response to selumetinib (AZD6244). Cancer research 2010;70(6):2264–73 doi 10.1158/0008-5472.CAN-09-1577.

57. Yaeger R, Mezzadra R, Sinopoli J, Bian Y, Marasco M, Kaplun E, et al. Molecular Characterization of Acquired Resistance to KRASG12C-EGFR Inhibition in Colorectal Cancer. Cancer discovery 2023;13(1):41–55 doi 10.1158/2159-8290.CD-22-0405.

58. Awad MM, Liu S, Rybkin, II, Arbour KC, Dilly J, Zhu VW, et al. Acquired Resistance to KRAS(G12C) Inhibition in Cancer. The New England journal of medicine 2021;384(25):2382–93 doi 10.1056/NEJMoa2105281.

59. Dilly J, Hoffman MT, Abbassi L, Li Z, Paradiso F, Parent BD, et al. Mechanisms of Resistance to Oncogenic KRAS Inhibition in Pancreatic Cancer. Cancer discovery 2024;14(11):2135–61 doi 10.1158/2159-8290.CD-24-0177.

60. Ebright RY, Dilly J, Shaw AT, Aguirre AJ. Response and Resistance to RAS Inhibition in Cancer. Cancer discovery 2025;15(7):1325–49 doi 10.1158/2159-8290.CD-25-0349.

61. King EA, Meyers M, Nomura DK. Induced proximity-based therapeutic modalities. Nature reviews Drug discovery 2026;25(3):175–203 doi 10.1038/s41573-025-01316-z.

