## Supplemental Figures for "Spatial modulation of RAF by RAF/MEK glue enables full-dose combination with pan-RAF inhibitor and potent RAS-mutant tumor-selective MAPK and growth inhibition"

### Figure S1 (related to Fig. 1)

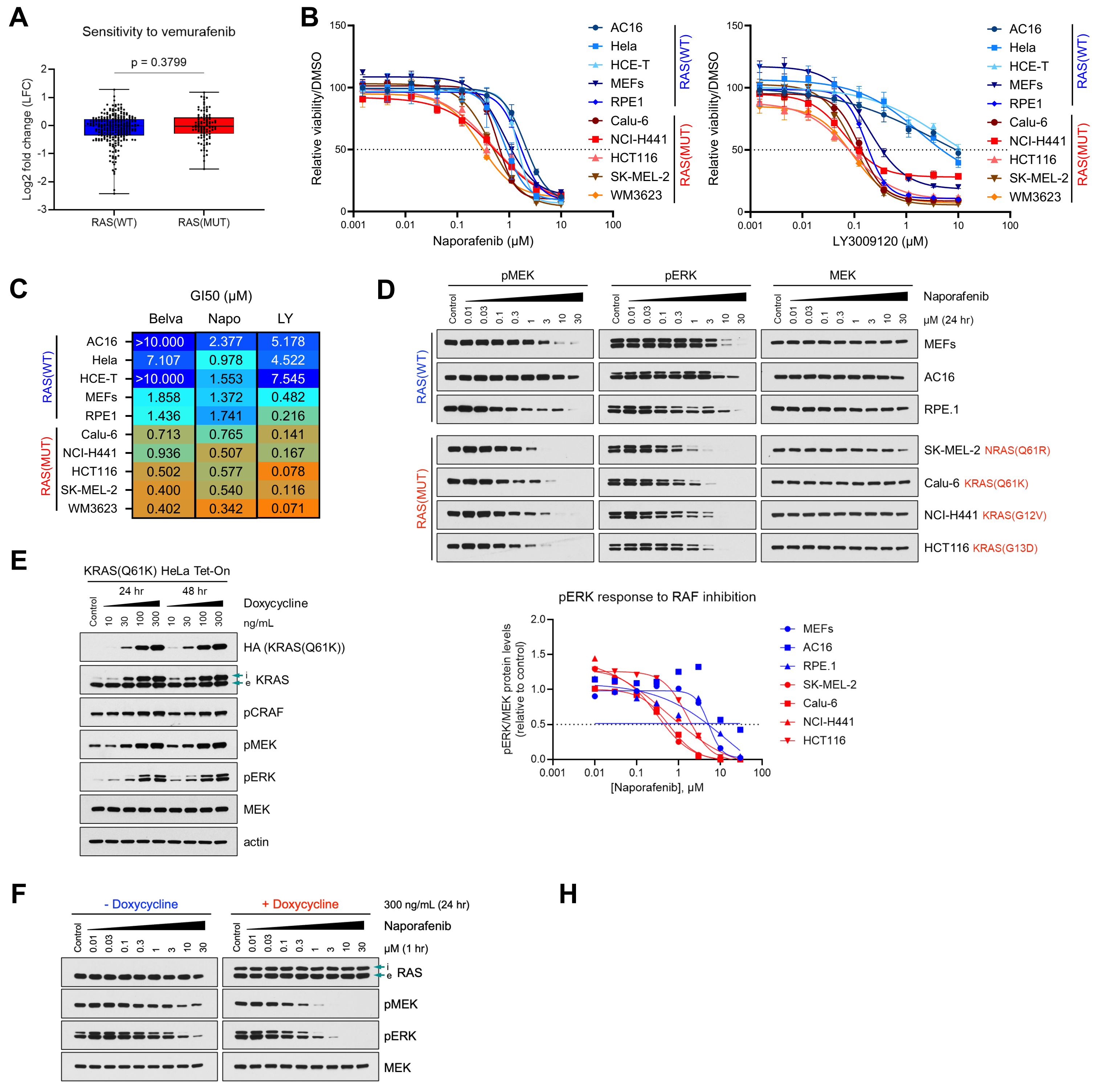


| | Belva | Napo | LY |
|---|---|---|---|
| AC16 | >10.000 | 2.377 | 5.178 |
| Hela | 7.107 | 0.978 | 4.522 |
| HCE-T | >10.000 | 1.553 | 7.545 |
| MEFs | 1.858 | 1.372 | 0.482 |
| RPE1 | 1.436 | 1.741 | 0.216 |
| Calu-6 | 0.713 | 0.765 | 0.141 |
| NCI-H441 | 0.936 | 0.507 | 0.167 |
| HCT116 | 0.502 | 0.577 | 0.078 |
| SK-MEL-2 | 0.400 | 0.540 | 0.116 |
| WM3623 | 0.402 | 0.342 | 0.071 |

H

G

pan-RAF-D

### Figure S1 (related to Fig. 1)

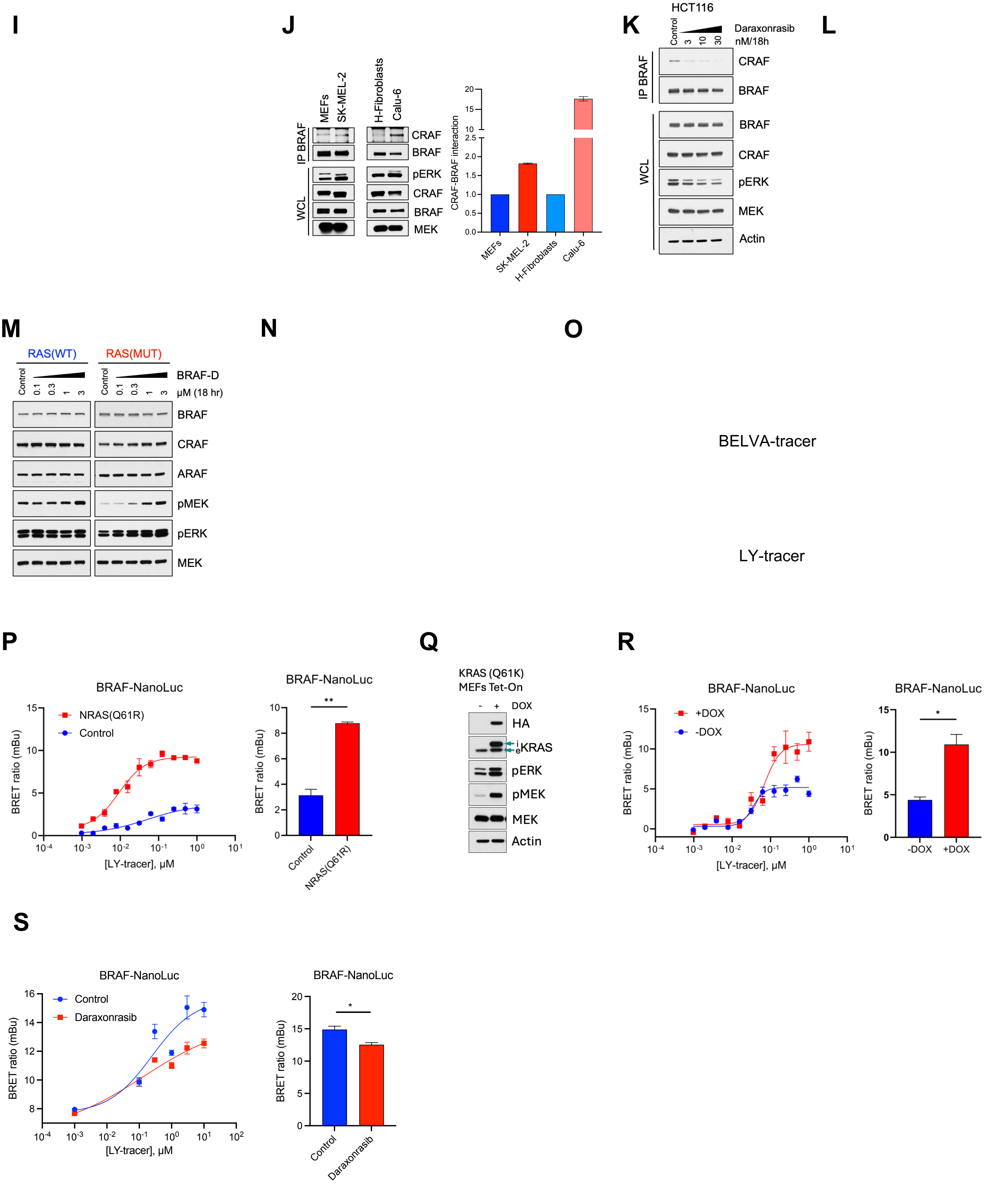

### Figure S2 (related to Fig. 2)

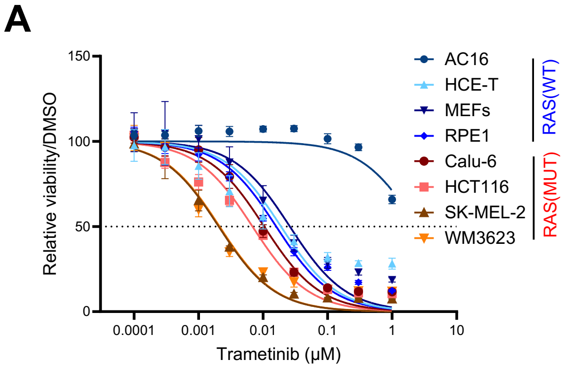


**B**

GI50 (μM)

| | | Cobi | Tram | Avuto |
|---|---|---|---|---|
| RAS(WT) | AC16 | 6.026 | >1.0000 | >10.000 |
| | HCE-T | 0.217 | 0.0190 | 1.660 |
| | MEFs | 0.046 | 0.0260 | 0.589 |
| | RPE1 | 0.094 | 0.0160 | 0.275 |
| RAS(MUT) | Calu-6 | 0.102 | 0.0100 | 0.544 |
| | HCT116 | 0.027 | 0.0068 | 0.146 |
| | SK-MEL-2 | 0.038 | 0.0020 | 0.075 |
| | WM3623 | 0.019 | 0.0021 | 0.054 |

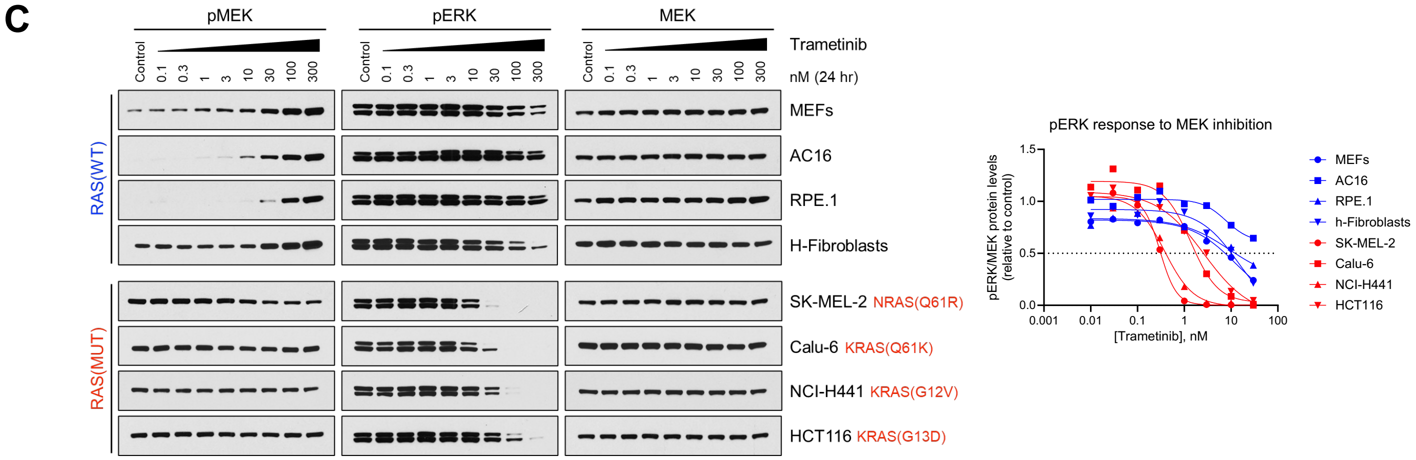


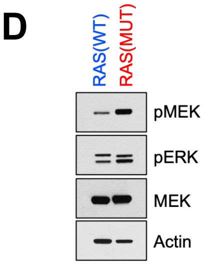

### Figure S3 (related to Fig. 3)

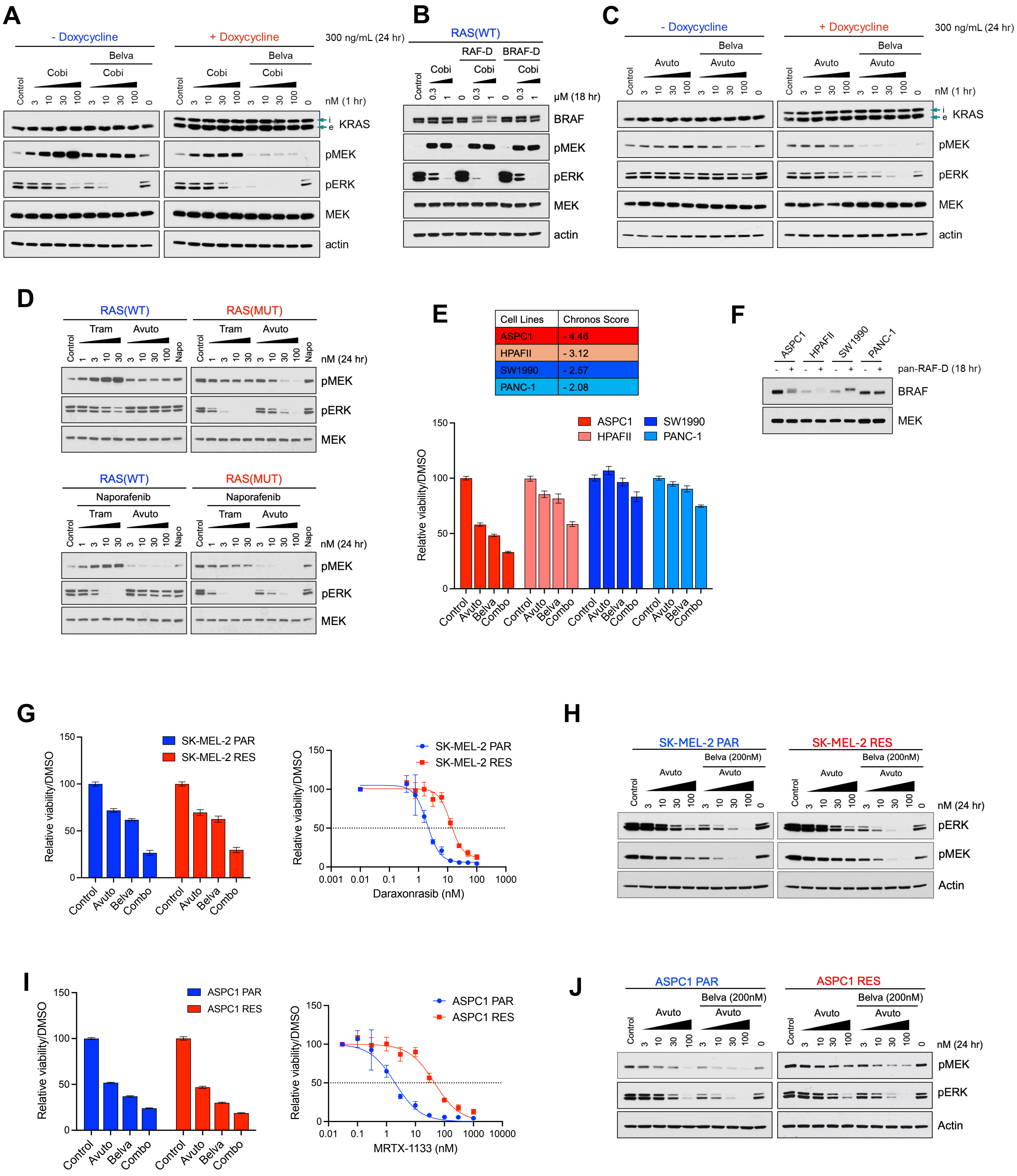

### Figure S4 (related to Fig. 4)

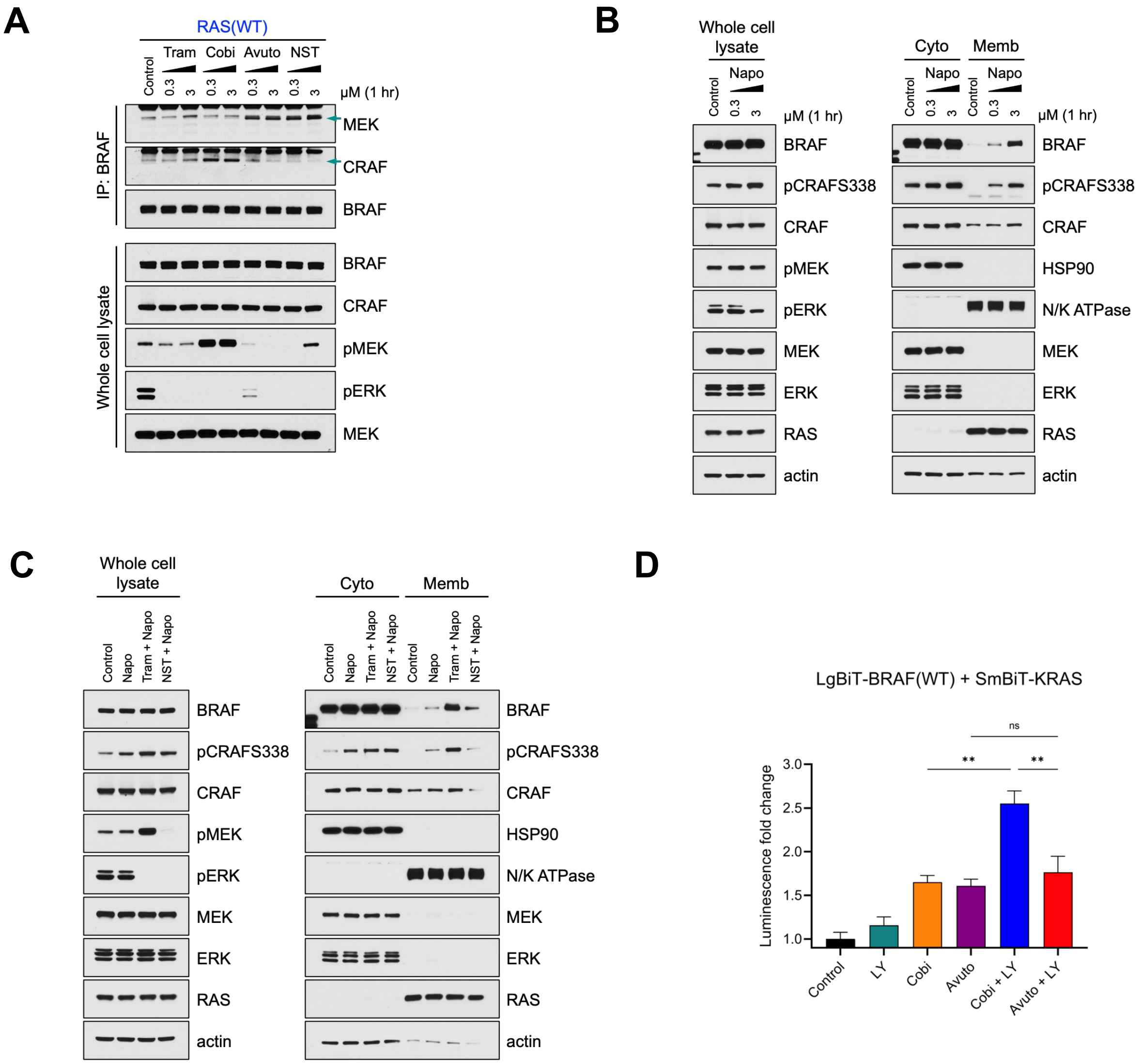

### Figure S5 (related to Fig. 5)

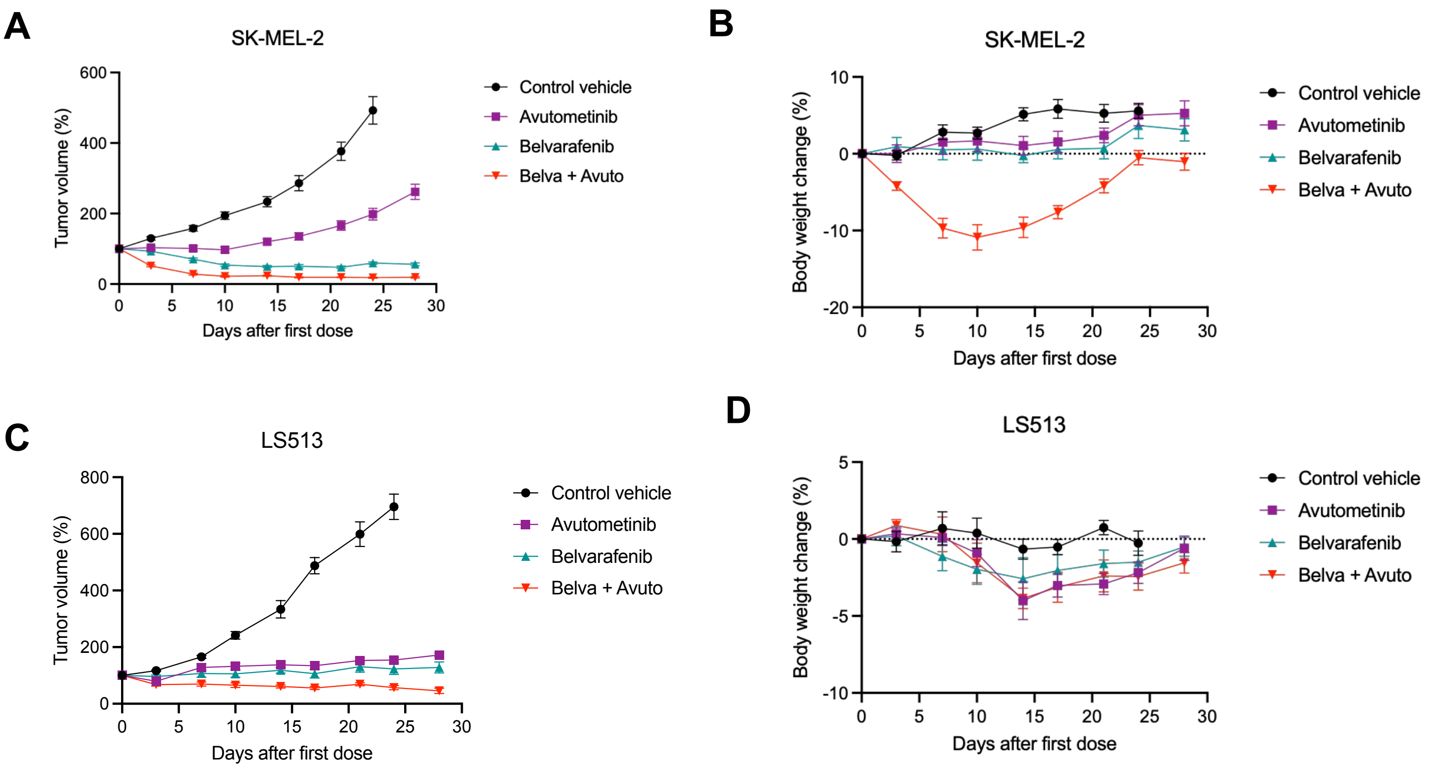

#### Figure S6 (related to Fig. 6)

A

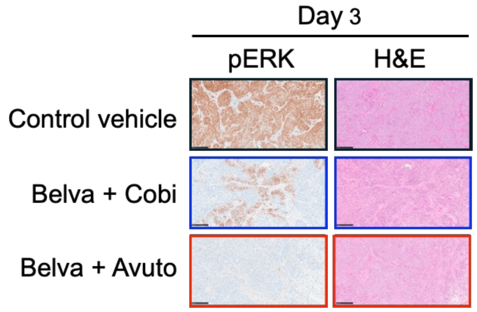


B

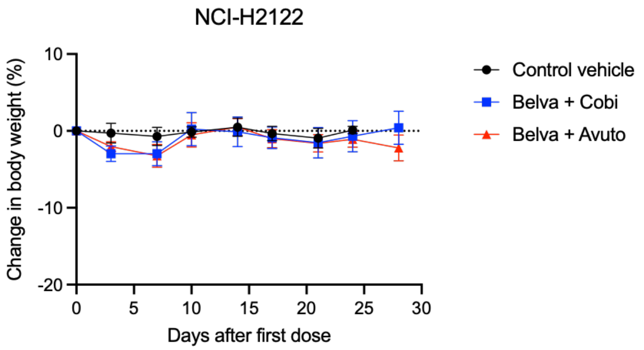
